# The Human Insula Encodes Somatotopic Representation of Motor Execution with an Effector-Specific Connectomic Map to Primary Motor Cortex

**DOI:** 10.1101/2025.05.19.653527

**Authors:** Panagiotis Kerezoudis, Michael A Jensen, Bryan T. Klassen, Gregory A. Worrell, Nick M. Gregg, Nuri F. Ince, Dora Hermes, Kai J Miller

## Abstract

Understanding motor representation in the human brain requires mapping beyond the primary motor cortex, into the distributed networks that coordinate complex movements. The insular cortex, a multifunctional hub buried within the Sylvian fissure, has been implicated in motor control through clinical observations and neuroimaging. Yet its precise relation to movement processing remains one of the least understood aspects of motor neurophysiology. To address this gap, we quantified electrophysiological changes from implanted depth electrodes in patients performing simple movement tasks combined with single-pulse electrical stimulation (SPES) to map functional connectivity. The movement data reveal distinct somatotopic representation bilaterally, as well as inter-effector regions that are active for different movement types. Hand representation is centered along the ventral aspect of the middle and posterior short gyri bilaterally, while tongue/mouth tuned sites cluster in the dorsal posterior short gyrus and the dorsal long gyri. Insular activity temporally follows the primary motor cortex (M1) and precedes movement onset. SPES revealed somatotopically-specific connectivity between corresponding sites in M1 and insula (hand-to-hand, tongue-to-tongue) and between bilateral insulae. These observations establish that somatotopy is a conserved property of distributed motor control incorporating the insula, with direct implications for basic and clinical neuroscience.

## INTRODUCTION

The topographic organization of neural function is a fundamental principle of brain architecture, wherein dedicated cortical regions process specific aspects of our interaction with the world. Such functional specialization manifests as somatotopy for body movements, retinotopy for visual space mapping, or tonotopy for auditory frequency processing.^1^ While somatotopic organization is well characterized on the precentral gyrus (primary motor cortex/M1), our understanding of movement representation in deeper brain regions remains less understood.^2^

The insular cortex, situated within the Sylvian fissure, is increasingly recognized as a hub integrating sensory, emotional, and cognitive information, with evidence suggesting an important role in motor control.^3,4^ Clinical data shows that up to 30% of patients with epilepsy exhibit motor deficits following insular procedures. These deficits are typically transient, with substantial functional recovery occurring within 1-2 weeks.^5,6^ Collectively, these observations suggest a nuanced, potentially integrative function of the insula in motor processing, yet the precise somatotopic arrangement within the human insula and its functional interplay with M1 remain inadequately defined.

Specifically, our understanding of insular movement representation has been limited by several key factors: its deep anatomical location that precludes access using traditional electrocorticographic grids and strips, limited temporal resolution of conventional neuroimaging techniques, and reliance on group-averaged data that obscures individual variability in functional organization. Stereo-electroencephalography (SEEG) recordings in human epilepsy patients provide a unique opportunity to overcome these limitations by enabling direct access to the insula with high-fidelity electrophysiology.^7,8^ Analysis of broadband spectral changes in SEEG voltage measurements allows quantification of local neuronal ensemble activity during precisely timed movement tasks, while single-pulse electrical stimulation (SPES) is a powerful tool for probing effective connectivity between motor cortical areas.^7,9^

Here, we leverage these SEEG tools to carefully delineate the somatotopic organization of movement representation in the human insula and elucidate its functional relationship with the primary motor cortex. This investigation provides critical insights into the role of the insula in motor control and advances our understanding of distributed motor networks in the human brain.

## RESULTS

Using a visually-cued, self-paced motor paradigm with intracranial recordings, we examined primary cortical and insular responses during hand, tongue & foot movement in 17 patients with epilepsy. Analysis of high-frequency broadband power (65-115 Hz) revealed clear somatotopic organization in insular activity. We observed distinct insular tongue movement representation in 15/17 subjects, while insular representation was captured for the hand and foot in 11 and 3 subjects, respectively **(Figure 1)**. Activation patterns were similar for left and right insulae. These tuned responses, though generally less pronounced than their primary motor cortex (M1) counterparts, showed distinct spatial clustering within our insular-specific coordinate system: hand sites concentrated along the ventral middle and posterior short gyri, while tongue sites predominantly occupied the dorsal posterior short and dorsal long gyri **(Figure 1)**. In addition to somatotopic representation, a shared-representation site, active during all movement types, was found in 4 subjects.

**Figure 1.**
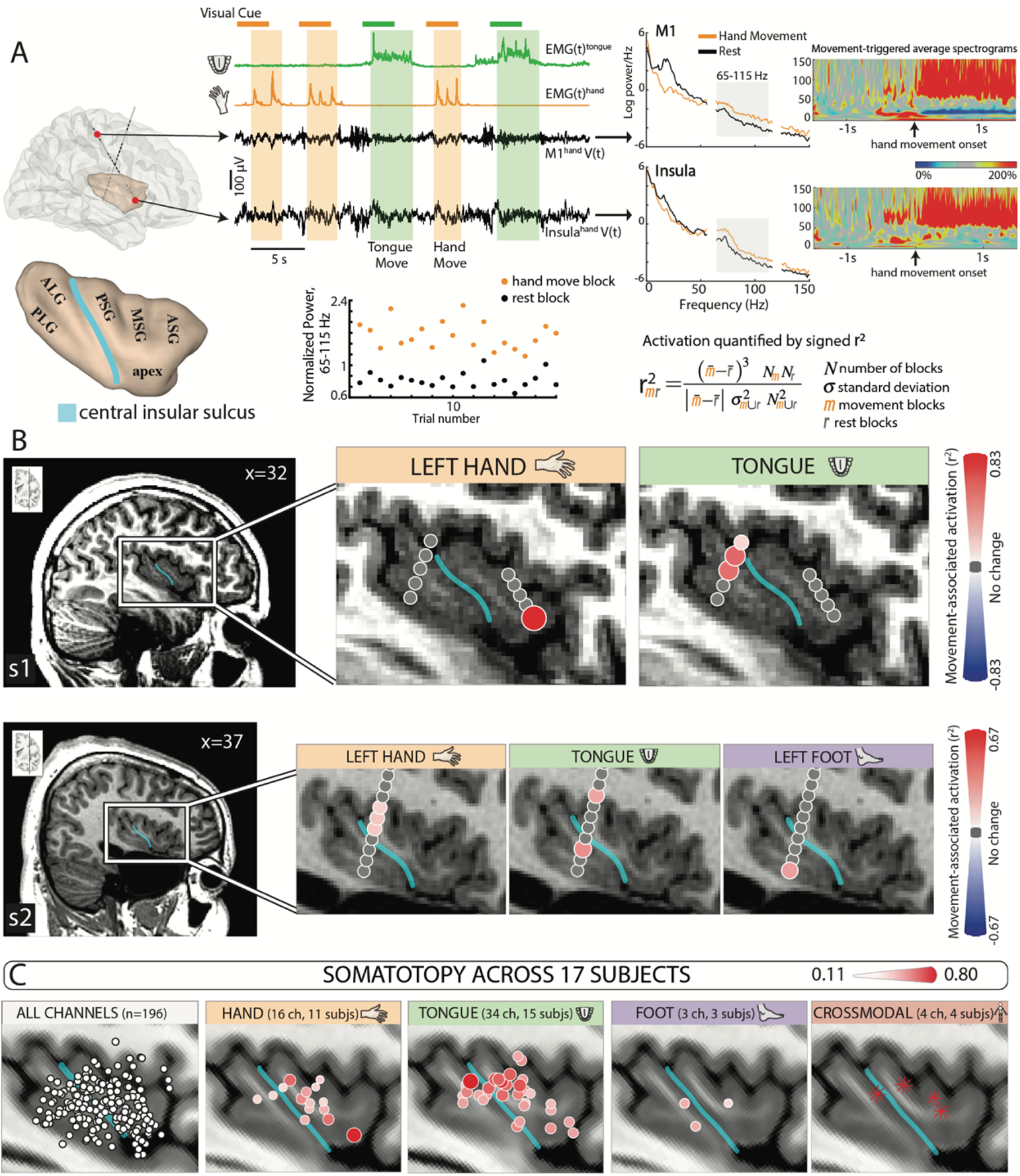
Insular movement representation is somatotopic. **(A)** The bipolar re-referenced signal for each stereo-EEG channel is divided into blocks of movement vs rest, the trial-average power spectrum in the 65-115 Hz range is extracted and the degree of movement activation is quantified using the signed-rank correlation coefficient (r^2^). The central sulcus of the insula (blue line) divides it into anterior and posterior regions. The anterior region consists of three short gyri (anterior/ASG, middle/MSG, posterior PSG) that converge at the insular apex, and the posterior region consists of two long gyri (anterior/ALG, posterior/PLG). **(B)** Activation r^2^ maps for hand (orange), tongue (green) and foot (purple) movement in the sagittal plane, for subject 1 (upper) and 2 (lower). Each map was scaled to the subject’s max r^2^ value. **(C)** Following transformation in insular stereotactic space (see Methods for details), all intra-insular channels (differential pairs) across all subjects are shown on the MNI-152 brain. The hand-tuned sites cluster along the middle and posterior short gyri bilaterally, while tongue-tuned sites clustered in the dorsal part of posterior short and anterior long gyri.

Temporal analysis in 9 patients with both M1 and insular somatotopic sites demonstrated that insular activation generally followed M1 activity during movement execution, with delays ranging from 41-266 ms for hand movement and 20-515 ms for tongue movements **(Figure 2)**. Both regions activated before movement onset (100% of M1 channels and 86% of insular channels), with M1 leading the sequence. We examined the connectivity of these motor-selective insular sites using SPES in two subjects, finding robust somatotopically-specific connectivity between corresponding M1 and insular sites, i.e. hand-to-hand and tongue-to-tongue **(Figure 3)**. In one patient with bilateral insular electrodes, we found reciprocal, somatotopically-specific connectivity between homologous insular regions. **(Figure 4)**.

**Figure 2.**
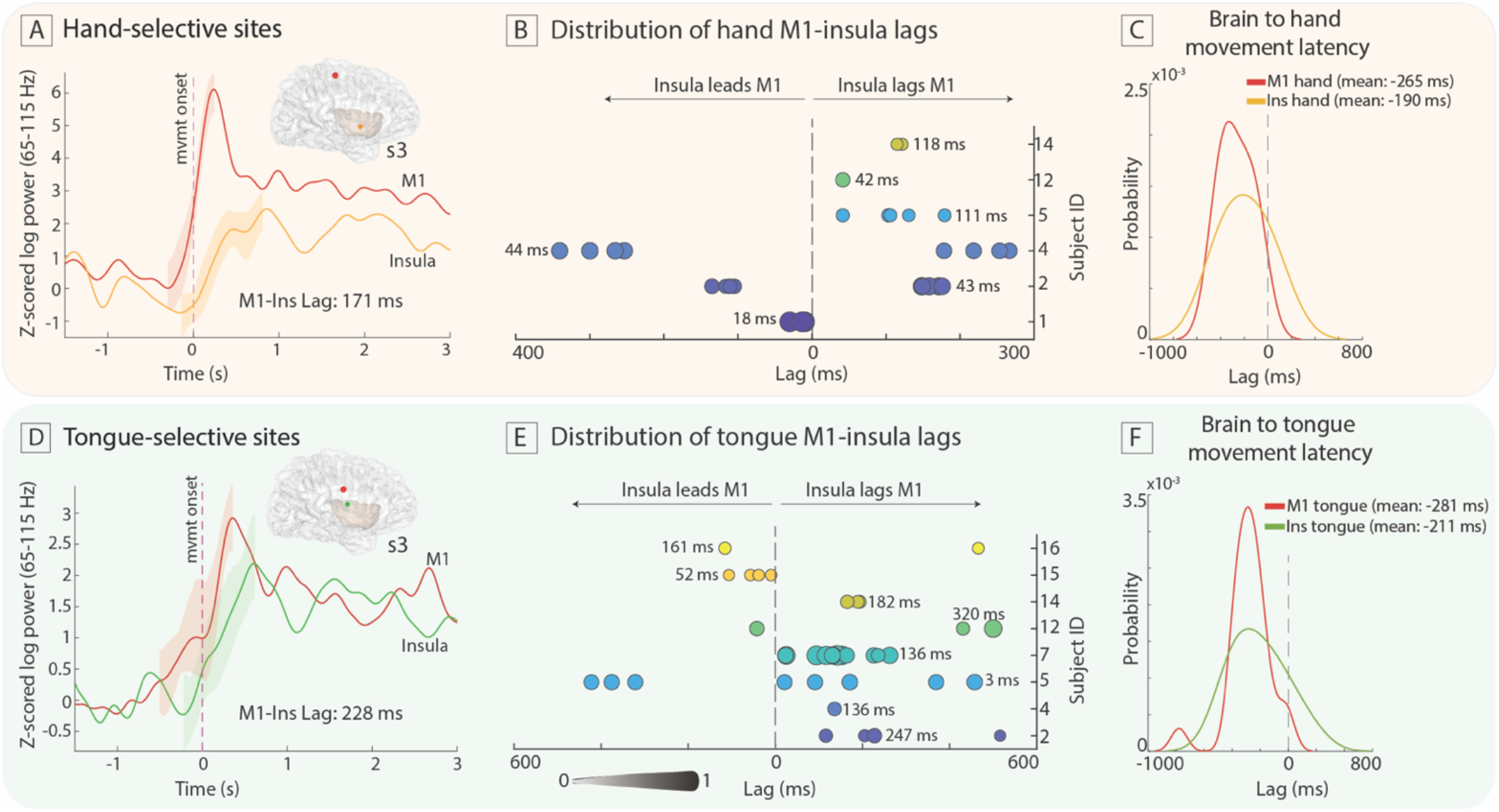
Insular activity generally follows M1 and precedes EMG during movement execution. **(A)** Demonstration of primary motor (M1), and insular hand movement-triggered average broadband power traces in subject 3, along with the standard error of the mean during the rising phase (see Methods for details). The M1 site preceded insular activity by 171 ms. **(B)** Distribution of hand M1-insular broadband power lags across 6 subjects. The numbers in each row correspond to the weighted average lag in each patient. Each circle is weighted by sqrt(r^2^_M1_*r^2^_ins_) and scaled to a maximum weight of 1. **(C)** Probability density functions of the distribution of M1 and insular channel to hand movement latencies. M1 and insular hand sites preceded EMG onset by 265 ms and 190 ms, respectively. **(D-F)** Similar temporal patterns emerged during tongue movement.

**Figure 3.**
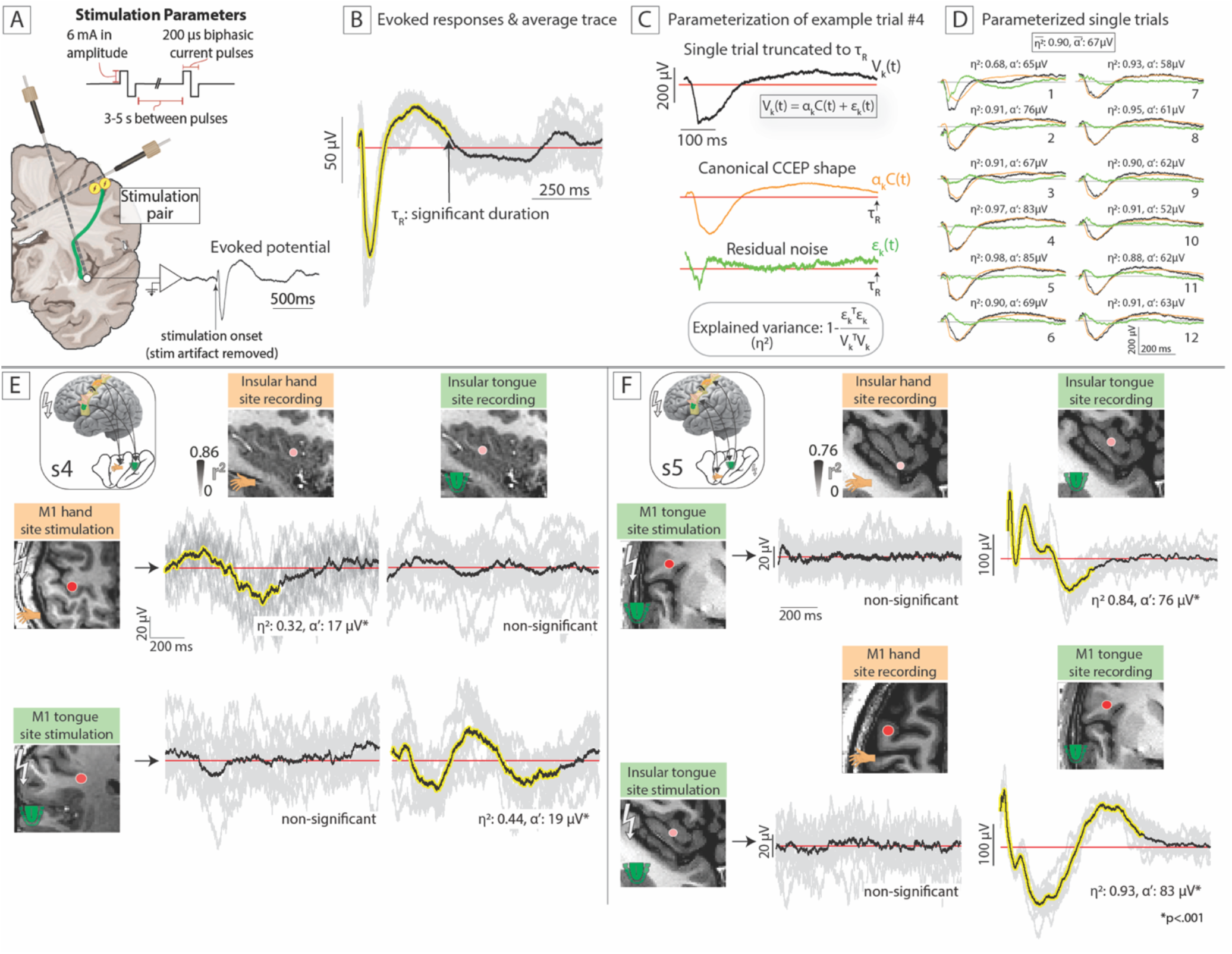
Somatotopic M1 sites exhibit effector-specific connections with corresponding insular sites. **(A)** Illustration of the CCEP experimental paradigm, showing a schematic of a brain (coronal slice) implanted with SEEG probes. Electrical stimulations were delivered at pre-specified channels and signals were recorded in all the other contacts. The stimulation parameters are shown in the box. **(B)** Summary of the Canonical Response Parameterization analytical approach for quantifying duration (τ_R_) and magnitude of significant evoked response. **(C)** Each stimulation trial is parameterized into a canonical response shape, defined by a_k_C(t), and residual noise ε_k_(t). **(D)** With this approach, we can extract the explained variance η^2^ and projection weight α′ for each trial as well as the average (α′=α/sqrt(time points)). **(E)** Single-pulse electrical stimulation (subject 4) at the somatotopically-tuned M1 hand and tongue sites showed significantly stronger evoked responses in the corresponding insular sites of the same type, but not the different type. **(F)** Similarly in subject 5, robust evoked responses were elicited upon stimulation of the M1 tongue site on the corresponding insular site and vice versa, but were not significant at the hand-selective sites.

**Figure 4.**
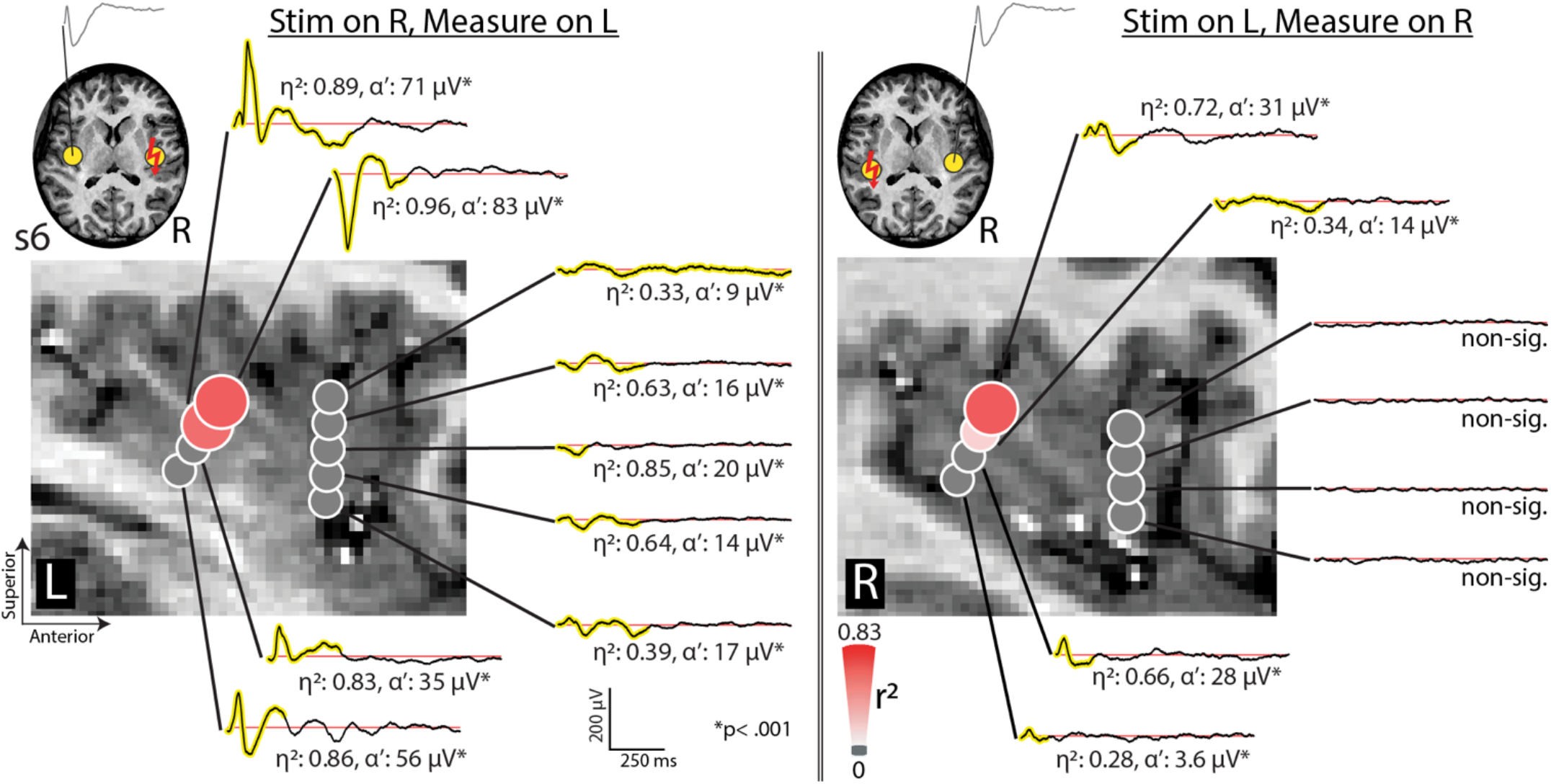
Reciprocal insular connections: somatotopic sites in the insula have robust connections with their counterpart in the contralateral side. Stimulation of the right tongue-tuned insular site (subject 6) in the posterior dorsal insula elicited robust responses in the left tongue-tuned site (same region), and vice versa.

## DISCUSSION

Our intracranial recordings reveal a robust, somatotopically-separated motor map in the human insula: generally hand-selective activity clusters along the ventral anterior cortex, whereas tongue-selective responses occupy dorsal posterior regions. Foot representation was sparsely detected, which may reflect evolutionary specialization of the insula for orofacial and manual dexterity control or could be a limitation of our spatial sampling. While the population-aggregate maps were overlapping, somatotopic separation was ubiquitous.

These electrophysiological findings significantly advance our understanding of insular motor representation, building upon earlier electrical stimulation and fMRI studies in non-human primates and humans that implicate the insula in motor control and tactile processing.^3,10–12^ The precise millisecond-scale latencies we document—where insular activity follows M1 but precedes movement onset—position the insula neither as a purely premotor region nor as a simple sensory processor. Instead, our data suggest the insula serves a dynamic role between action execution and sensory perception. This arrangement allows the insula to contextualize ongoing movements within a broader sensorimotor framework, potentially facilitating the distinction between self-generated and external sensory experiences during movement.^4,13^

Connectivity patterns seem to preserve ordered somatotopic representations. Previous research using tracing studies in macaques, diffusion MRI tractography in humans, and cortico-cortical evoked potentials have established structural connectivity between the insula and the pre-/post-central gyri.^14–17^ Our results extend this work by demonstrating that these connections are specifically organized according to somatotopic maps. When stimulating hand-selective sites in M1, we observed responses primarily in hand-selective insular regions. Similarly, stimulation of tongue-selective M1 sites activated corresponding tongue-selective insular regions, and vice-versa.

This representation-preserving connectivity architecture supports a modular organization of movement processing networks. The observed topology enables parallel processing of effector-specific information rather than a fully distributed, intermingled body representation within each anatomic structure. Specifically, the precise wiring along with the observed temporal dynamics may position the insula ideally to process efference copy signals. In such a model, signals from the M1 cortex to association areas serve as internal predictive models of expected sensory feedback.^18^ These predictive mechanisms are essential for distinguishing self-generated from external stimuli and for fine-tuning ongoing movements through sensory prediction error. More complex behavioral paradigms may further examine the role of the insula, if any, in processing efference copies.

Deciphering the functional anatomy and connectivity of the human insula has important clinical applications, particularly for epilepsy treatment. Insular epilepsies exhibit distinct seizure sensorimotor symptoms that correlate with our mapped regions^19^: seizures originating in the posterior insula frequently begin with laryngo-oro-facial motor automatisms, throat constriction, or unilateral paresthesias/tonic contractions of the upper extremity before propagating to the perirolandic cortex.^20^ Our discoveries can therefore inform epilepsy teams in refining SEEG implantation and neurostimulation strategies based on specific seizure semiology. Beyond epilepsy, the precise localization of hand and tongue representations offers a roadmap for developing targeted neuromodulation therapies for speech disorders, dysphagia, and hand-related motor dysfunctions. Individualized functional mapping will be crucial for optimally personalizing therapeutic outcomes rather than relying on group-averaged maps that obscure discrete functional zones within subjects.

In summary, we demonstrate a networked motor homunculus in the human insula with somatotopic organization and specific connectivity patterns, opening new avenues for both basic and clinical neuroscience.

## MATERIALS AND METHODS

### Ethics statement

The study was conducted according to the guidelines of the Declaration of Helsinki and approved by the Institutional Review Board of the Mayo Clinic (IRB number 15-006530), which also authorizes sharing of de-identified data. Each patient or parental guardian provided informed consent as approved by the IRB. All T1 MRI sequences were de-faced prior to publication using an established technique, to avoid potential identification.^1^

### Participants

A total of 17 patients (9 females) participated in the current study, aged 12-37 (n=6 ≽ 18 y/o) with medically refractory epilepsy undergoing intracranial recordings with SEEG leads as part of their seizure workup. All but three patients (s2, s4, s5) had left-sided dominance for language. A mean of 14 SEEG leads (range 10-17) per patient were placed. Electrode locations were planned by the clinical epilepsy team based on clinical semiology, scalp EEG studies, and brain imaging.^2^ No plans were modified to accommodate research, nor were extra electrodes placed. All experiments were performed in the epilepsy monitoring unit or the pediatric intensive care unit. A summary of patient demographics and other clinical details is provided in **Supplemental Table 1**.

### Lead implantation

The circumferential platinum depth electrode contacts were 0.8 mm in diameter and 2 mm in length **[Supplemental Figure 1]**. Each lead contained 10-18 electrode contacts. Surgical targeting and implantation were performed in the standard clinical fashion. Intraoperatively, anchoring bolts were placed into stereotactically aligned 2.3 mm holes in the skull and then leads were advanced to target through the bolts.^2^ The insular cortex was typically targeted by two focused leads entering from oblique trajectories in the parasagittal plane, with one directed towards the anterior insula from a posterior trajectory, and one to the posterior insula from an anterior trajectory. However, any contacts from other leads targeting different structures that passed through the insular cortex were also included. Left hemispheric insular leads were placed in 7 patients, right hemispheric in 8 patients, while 2 patients were implanted with bilateral insular leads (s6, s10). A third electrode was placed in the middle insula in 2 subjects (s2, s12) based on clinical indication.

### Electrode localization and re-referencing

Electrode-to-anatomic localizations were determined by co-registering the post-implant CT scan to the pre-implant MRI. Each preoperative T1 MRI was realigned to the anterior/posterior commissure stereotactic space (AC-centered), and then co-registered to the post-implant CT using SPM12.^3^

All data were bipolar re-referenced, highpassed (3^rd^ order, zero-phase Butterworth filter, stopband 0.05 Hz, passband: 0.5 Hz, passband ripple 3dB) and notch-filtered for 60Hz line noise and its harmonics (4^th^ order, zero-phase Butterworth filter). Channels that were noisy, located in the seizure onset zone or contaminated with significant noise/interictal discharges were excluded from further analysis. Finally, we carefully inspected the location of all channels on co-registered CT/MRI to a) exclude contacts labeled as insular that were very close to putamen or operculum (n=15) and b) include insular contacts from non-insular focused leads (n=10). Across the 17 subjects, a total of 39 insular focused leads (left side: 18; right side: 21) and 196 bipolar channels (left side: 89; right side: 107) were analyzed, with a mean of 10 channels per patient (range: 6– 24).

### Behavioral motor paradigm

Tasks were performed at least 24 hours after surgery to allow for settling of brain irritation and artifact generation secondary to the implantation process. Subjects were asked to perform three types of visually-cued, self-paced (∼1 Hz) movements, and to remain still during interleaved rest periods (blank screen): 1) opening and closing of the contralateral hand to the hemisphere of SEEG montage, 2) side-to-side movement of the tongue with mouth closed, and 3) alternating dorsi-and plantar flexion of the contralateral foot. A total of twenty cues (trials) of each movement type were shuffled in random order and move/rest trial cues were each presented for 3 seconds **[Supplemental Figure 1]**. This task paradigm was chosen based upon prior work, which has produced clear and reliable primary sensorimotor mapping results in SEEG recordings.^4,5^ Stimuli were presented on a 53 × 33 cm screen, 80-100 cm from the subject’s face. If subjects did not stay engaged with the task, the experimental run would be stopped and re-run later. Muscle movement was captured with electromyography (EMG) using surface pads placed on forearm flexors/extensors (hand), base of chin (tongue), and anterior tibialis (foot).^4^ In four subjects (s2, s4, s8, s9), epoch segmentation for tongue movement was based on screen cue due to suboptimal EMG signal. All SEEG and EMG signals were measured in parallel at 1200Hz (g.HIAmp system, gTec, Austria) and were synchronized with screen cues using the BCI2000 software.^6^ Data were segmented into movement and rest periods using synchronized EMG as previously described.^4^

### Power spectral snapshots

Averaged power spectral densities (PSDs) for each movement trial were calculated from 1 to 150 Hz in 1 Hz frequency bins using Welch’s averaged periodogram method, with 0.5 s Hann windows to attenuate edge effects, and 0.25 s overlap. The averaged PSD for each movement or rest trial was normalized to the global mean across the entire experimental run for each channel. This was performed, because brain signals of this type generally follow a 1/f power law shape and therefore lower frequency features dominate in the absence of normalization.^7^ From each of these normalized single trial PSDs, averaged power in a broadband high frequency band (65-115 Hz) was extracted for subsequent analysis.^4^ This band was chosen because it captures broadband activity above the known range of most oscillations that might contaminate it, and avoids ambient line noise at 60 Hz and 120 Hz. It should be noted that we did not observe significant modulation in the canonical low-frequency oscillation range (e.g. mu or beta) in the insular channels during movement vs rest (though they were observed in M1) **[Supplemental Figure 2]**, and therefore oscillations were not explored further in the present study.

### Quantification of activation during movement vs rest

For each bipolar re-referenced channel, we calculated separate signed r^2^ cross-correlation values of the mean spectra from 65-115 Hz for each movement modality. Each channel’s r^2^ value was determined by comparing mean power spectra between each movement trial and rest. We only compared movement trials with rest periods that followed that same movement type, in order to minimize the cross effects of movement-specific rebound effects. The formula for calculating the signed cross-correlation r^2^ coefficient is the following:

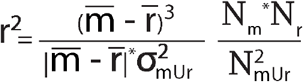

where m denotes power samples from movement, r denotes samples from rest and the overline (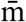 and 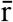) denotes sample mean. m∪r represents combined movement and rest power sample distributions. N_m_ and N_r_ denote the total number of movement and rest samples and N_mUr_=N_m_+N_r_ denotes the aggregate of both. To calculate a p-value for each channel and each movement type, we performed an unpaired two-sample t-test comparing broadband power for movement vs rest trials. We subsequently plotted the r^2^ values for each bipolar re-referenced channel on coronal, sagittal and axial MRI slices (see section below on *Electrode visualization*). This analytical approach is graphically summarized in **Figure 1**.

### Electrode visualization

Plotted points for brain activity represent an interpolated midpoint between the two electrodes that make up each differential pair channel. In each activity map, channels were plotted using the SEEGVIEW package, which slices at fixed intervals and projects lead contacts to the closest selected slice.^8^ The goal of this tool is to visualize results of analysis in an interpretable and clinically familiar manner. Given projection distance increases alongside slice thickness, channel reflecting activity *actually* in the gray matter may *artifactually* appear to be in white matter following projection to the nearest slice. The size and color intensity of the circle for each channel were co-scaled to the ‘global maximum’ (the maximum r^2^ value across all three movement types for each patient) and all insignificant channels were plotted with a grey circle of fixed small diameter.

### Somatotopic and shared representation

Channels were classified as somatotopically-tuned for a specific effector (hand, tongue, or foot) when they met two criteria: they showed a significant activation (r^2^ > 0.10, p < 0.05) for that particular movement and no significant activation for the other movements. In contrast, inter-effector insular channels with shared representation were identified based on overlapping movement responses, requiring significant correlations (r^2^ > 0.10, p < 0.05) for all three movement types (hand, tongue, and foot).

### Time-frequency spectrograms

We generated time-frequency approximations after convolving the voltage time-series data with a Morlet wavelet (5 cycles) in order to estimate the amplitude and phase of the signal across the frequency range of 1-150 Hz for every point in time. Time-frequency analysis was performed in a window 1.5 s before to 1.5 s after the onset of each movement trial (or screen cue in cases of suboptimal EMG, as discussed above), divided by the average power during a period of -1.5 s to -0.5 s prior to movement onset, and then averaged across all movement trials of the same type to produce a movement-triggered average dynamic spectrum.

### Baselining and averaging the neural signal

The issue of baselining the broadband signal is somewhat nuanced. Baseline periods may be timed to the cue onset or to the movement onset. If timed to cue onset, one typically picks a period of 500 to 1000 ms ending at the time of cue onset. If timed to movement onset, the baseline is typically set to begin at ∼1500-1000 ms prior to movement onset, and end at ∼300-500 ms prior to movement onset. Baselining may be performed on a trial by trial basis, or globally (see **Supplemental Figure 4** for the different combinations). In the present study, we chose a global baseline to minimize bias, by normalizing the entire broadband trace with respect to a pre-stimulus distribution that is aggregated across all stimulus types and trials (−500 ms to 0 s prior to cue onset). This approach, while maximally general, risks missing an effect if there is a drifting or a dynamic brain state, the variation of which would be lost by picking a global distribution for baseline.

Consideration of the issue of triggered times for averaging across trials can be useful in determining to what degree a response is stimulus-locked or movement-locked. We initially performed both, the results of which are shown in **Supplemental Figure 5**. The coefficient of variation (defined as standard deviation/mean) during the first 500 ms following screen cue and movement onset for somatotopically-tuned channels showed that both M1 and insular dynamics were movement-locked rather than stimulus-locked. As a control, we show that visual areas of the brain are stimulus-locked, as expected. Therefore, we decided to use movement onset-locked analysis for the broadband signal.

### Broadband power time series

We obtained broadband power time series for each channel by (1) band-passing the channel voltage with a third-order Butterworth filter in 10 Hz bands between 65 and 115 Hz, (2) applying the Hilbert transform, (3) squaring each 10 Hz time series and (3) adding the 10 Hz time series together. The resulting signal was subsequently log-transformed, smoothed with a 500 ms Gaussian kernel, and z-scored based on a 500 ms window prior to all screen cues for movement (i.e. hand, tongue or foot) to obtain a broadband power timeseries. Each power timeseries was averaged over movement trials to independently generate hand-triggered, tongue-triggered and foot-triggered traces that were each used for latency/lag analysis (see section below). We also generated 95% confidence intervals for each average trace using a 1000-sample bootstrapping resampling approach. The decision to use log-transformed data for analysis was based on the fact that high frequency broadband power follows a log-normal distribution, which allows for Gaussian statistics.^9^ This is shown for the present data in **Supplemental Figure 6**.

### Lag between somatotopic M1 and insular channels

We quantified the temporal relationship between somatotopic primary motor (M1) and insular channel activation patterns using a novel rising phase isolation approach (see **Supplemental Table 2** for somatotopic sites per subject). This method begins by calculating the first derivative of each input signal (i.e M1 and insula movement-triggered averages) to identify positive rate changes, while setting negative changes to zero. The algorithm then detects continuous rising phases by identifying transitions between rising and falling segments. To enhance detection reliability, the method incorporates a consolidation protocol that merges rising phases separated by intervals shorter than 200ms. The analysis specifically targets the rising phase segment containing the global maximum amplitude, enabling isolation of the predominant activation pattern while excluding minor transient fluctuations. The temporal offset between M1 and insular activation is then determined by computing the sliding dot product cross-correlation between the corresponding segments, with the time point corresponding to the global maximum of the cross-correlation indicating the activation lag between the two regions. A detailed visualization of this methodological approach is provided in **Supplemental Figure 7**.

We also quantified the effect of the Gaussian window smoothing on this statistic by ranging its length from 100 to 1000 ms when extracting the broadband power time series (see **Supplemental Figure 7C**). It can be appreciated that by enlarging the window length, the lag progressively converged to an asymptotic value. In our data, we found that this convergence to the asymptote value happened by 500 ms in all cases examined.

### Latency of brain activity with respect to movement onset

While there are many potential ways to define the onset time of brain activity, to assess whether brain “activation” precedes movement, we feel that the most robust one is the midpoint between the initiation of the rising phase of brain activity (as defined in the methods section above), and the inflection point of the rising phase (when the second derivative is zero). These two points represent the most and least conservative estimates (see **Supplemental Figure 8**). We found that somatotopic insular activity preceded EMG in all but 18/21 channels, and in 43/43 channels for M1 somatotopic sites.

### Group-level analyses

The intrinsic person-to-person variability in insular anatomy poses significant challenges for traditional neuroimaging registration approaches that rely on non-linear transformation of individual brains into a common space, such as MNI-152. Such transformations, while widely used in neuroimaging research, have several important limitations, including failing to capture the significant inter-individual variability in brain structure and morphology, registration errors in region with high variability or presence of pathology and smoothing effect during the warping process obscuring fine structural details.^10^ Notably, they do not address the relative internal rotations of insular gyri compared to the standard direction of stereotactic spaces.

To address these limitations and preserve individual anatomical characteristics in the insular region, we developed a novel approach using an insular coordinate system. Our method begins with the affine transformation of individual patient brain MRI into the standard AC-PC space (origin at the mid-commissural point). This initial step provides a standardized orientation for all brain images and serves as a common starting point for further analysis. Following the AC-PC alignment, we subsequently performed a transformation into an insular-specific space.

This transformation is based on five anatomical landmarks identified on each patient’s MRI: four points along the central sulcus of the insula that define the primary insular axis and one inferior point where the middle cerebral artery (MCA) typically bifurcates/trifurcates, which serves as the anterior-posterior extent marker.

To transform into insular space, we calculate two angles: θ (theta) and φ (phi) (**Supplemental Figure 9**). The angle θ represents the rotation in the axial plane between the AC-PC line and the insular axis when viewed from above. The angle φ represents the rotation in the sagittal plane between the AC-PC line and the insular axis when viewed from the side. The brain volume then undergoes a series of rotations using these angles: First, a rotation through the angle θ is performed in the axial plane, followed by a rotation through the angle φ in the sagittal plane. Finally, a translation is applied to position the coordinate system origin along the insular axis, with the anterior-posterior origin at the MCA turning point. A demonstration of this process in our custom-built graphical user interface in MATLAB (Mathworks, USA) is shown in **Supplemental Figure 10**. The same transformation was applied to MNI-152 (2009c nonlinear asymmetric, non-skull stripped) brain volume and somatotopic weights from channels across all patients were plotted in this rotated brain to illustrate group-level information.

### Brain-stimulation evoked potentials (BSEPs): data collection and analysis

#### Stimulation parameters

Selected electrode pairs in subjects 4, 5 and 6 were stimulated 12 times (trials), each with a single biphasic pulse of 200 μs duration and 6 mA amplitude every ∼5 seconds using a g.tec g.Estim PRO electrical stimulator (g.tec, Schiedlberg, Austria) **(Figure 3)**.^11,12^ A licensed neurologist was present during each stimulation experiment and seizure rescue medications were available in the unlikely event of seizure onset (although this has not happened in our research program, likely due to the very small amount of charge density involved).

#### Pre-processing of stimulation data

Stimulation sites that overlapped seizure onset zones, according to physician records, were excluded from analysis. Furthermore, electrodes and individual trials were then visually inspected, and those that contained artifacts or epileptiform activity. All data had at least 10 remaining stimulation trials. Data in all subjects were bipolar re-referenced, high-pass filtered above 0.5 Hz with a second-order Butterworth filter and notch-filtered for 60 Hz line noise (and its harmonics), as mentioned earlier for the behavioral analysis. To avoid artifact caused by electrical stimulation or volume conduction (largest in the first 1-8 ms), we focused our analysis on responses after 15ms and excluded artifact-contaminated electrodes in the same lead.^13–15^

#### Canonical response parameterization (CRP)

The CRP approach allows for parameterization of evoked potentials from single-pulse electrical stimulation (SPES) using a naive, data-driven technique. For detailed description, we refer the reader to an original article explaining the methodology as well as a subsequent publication applying the technique to characterize distinct electrophysiological signatures in the human limbic network.^11,16^ Instead of relying on predetermined shapes or polarity of evoked responses (e.g. N1 and N2 peaks), CRP discovers structure empirically, allowing for comparison among different brain areas with dissimilar trace shapes.

The CRP method characterizes each stimulation response in the form of: V_k_(t) = a_k_C(t) + ε_k_(t). Each individual trial *k* is represented as a projection of a canonical BSEP form *C(t)*, scaled by a scalar *α*_*k*_, with residual *ε(t)*. This way, we can analyze the evoked responses between somatotopic M1 and insular sites and quantify the mean projection weight (*α*′, normalized by the square root of number of time points to obtain magnitude in μV), and the mean explained variance (η^2^) for each stimulation-recording channel pair.

## Data availability

All anonymized data recorded necessary to interpret, verify and extend the research will be made publicly available at Open Science Foundation.

## Code availability

All code necessary to reproduce findings will be made publicly available on GitHub following the publication of the manuscript. All analysis was conducted in MATLAB (R2022b, Mathworks, Natick, MA, USA). Figures were produced in MATLAB and finally edited in Adobe Illustrator 2024 (Adobe, San Jose, CA, USA).

## References

1. Patel, G. H., Kaplan, D. M. & Snyder, L. H. Topographic organization in the brain: searching for general principles. Trends Cogn. Sci. 18, 351–363 (2014).

2. Meier, J. D., Aflalo, T. N., Kastner, S. & Graziano, M. S. A. Complex organization of human primary motor cortex: a high-resolution fMRI study. J. Neurophysiol. 100, 1800–1812 (2008).

3. Kurth, F., Zilles, K., Fox, P. T., Laird, A. R. & Eickhoff, S. B. A link between the systems: functional differentiation and integration within the human insula revealed by meta-analysis. Brain Struct. Funct. 214, 519–534 (2010).

4. Molnar-Szakacs, I. & Uddin, L. Q. Anterior insula as a gatekeeper of executive control. Neurosci. Biobehav. Rev. 139, 104736 (2022).

5. Mullatti, N. et al. Stereotactic thermocoagulation for insular epilepsy: Lessons from successes and failures. Epilepsia 60, 1565–1579 (2019).

6. Kerezoudis, P. et al. Insular epilepsy surgery: lessons learned from institutional review and patient-level meta-analysis. J. Neurosurg. 1–13 (2021).

7. Jensen, M. A. et al. A motor association area in the depths of the central sulcus. Nat. Neurosci. 26, 1165–1169 (2023).

8. Miller, K. J. & Fine, A. L. Decision-making in stereotactic epilepsy surgery. Epilepsia 63, 2782–2801 (2022).

9. Huang, H. et al. Electrical Stimulation of Temporal and Limbic Circuitry Produces Distinct Responses in Human Ventral Temporal Cortex. J. Neurosci. 43, 4434–4447 (2023).

10. Afif, A., Minotti, L., Kahane, P. & Hoffmann, D. Anatomofunctional organization of the insular cortex: a study using intracerebral electrical stimulation in epileptic patients. Epilepsia 51, 2305– 2315 (2010).

11. Mutschler, I. et al. Functional organization of the human anterior insular cortex and its relation to the amygdala. Neuroimage 47, S106 (2009).

12. Jezzini, A., Caruana, F., Stoianov, I., Gallese, V. & Rizzolatti, G. Functional organization of the insula and inner perisylvian regions. Proc. Natl. Acad. Sci. U. S. A. 109, 10077–10082 (2012).

13. Dosenbach, N. U. F., Raichle, M. E. & Gordon, E. M. The brain’s action-mode network. Nat. Rev. Neurosci. 26, 158–168 (2025).

14. Almashaikhi, T. et al. Functional connectivity of insular efferences. Hum. Brain Mapp. 35, 5279– 5294 (2014).

15. Catani, M. et al. Short frontal lobe connections of the human brain. Cortex 48, 273–291 (2012).

16. Felleman, D. J. & Van Essen, D. C. Distributed hierarchical processing in the primate cerebral cortex. Cereb. Cortex 1, 1–47 (1991).

17. Groenendijk, I. M. et al. Whole brain 7T-fMRI during pelvic floor muscle contraction in male subjects. Neurourol. Urodyn. 39, 382–392 (2020).

18. Wolpert, D. M. & Ghahramani, Z. Computational principles of movement neuroscience. Nat. Neurosci. 3 Suppl, 1212–1217 (2000).

19. Singh, R. et al. Mapping the insula with stereo-electroencephalography: The emergence of semiology in insula lobe seizures. Ann. Neurol. 88, 477–488 (2020).

20. Jobst, B. C. et al. The Insula and Its Epilepsies. Epilepsy Curr. 19, 11–21 (2019).

## References

1. Bischoff-Grethe, A. et al. A technique for the deidentification of structural brain MR images. Hum. Brain Mapp. 28, 892–903 (2007).

2. Miller, K. J. & Fine, A. L. Decision-making in stereotactic epilepsy surgery. Epilepsia 63, 2782–2801 (2022).

3. Ashburner, J. et al. SPM12 manual. Wellcome Trust Centre for Neuroimaging, London, UK 2464, (2014).

4. Jensen, M. A. et al. A motor association area in the depths of the central sulcus. Nat. Neurosci. 26, 1165–1169 (2023).

5. Jensen, M. A. et al. Functional Mapping of Movement and Speech Using Task-Based Electrophysiological Changes in Stereoelectroencephalography. bioRxiv (2024) doi:10.1101/2024.02.29.582865.

6. Schalk, G., McFarland, D. J., Hinterberger, T., Birbaumer, N. & Wolpaw, J. R. BCI2000: a general-purpose brain-computer interface (BCI) system. IEEE Trans. Biomed. Eng. 51, 1034–1043 (2004).

7. Miller, K. J., Sorensen, L. B., Ojemann, J. G. & den Nijs, M. Power-law scaling in the brain surface electric potential. PLoS Comput. Biol. 5, e1000609 (2009).

8. Huang, H., Valencia, G. O., Hermes, D. & Miller, K. J. A canonical visualization tool for SEEG electrodes. Conf. Proc. IEEE Eng. Med. Biol. Soc. 2021, 6175–6178 (2021).

9. Miller, K. J. et al. Human motor cortical activity is selectively phase-entrained on underlying rhythms. PLoS Comput. Biol. 8, e1002655 (2012).

10. Evans, A. C., Janke, A. L., Collins, D. L. & Baillet, S. Brain templates and atlases. Neuroimage 62, 911–922 (2012).

11. Ojeda Valencia, G. et al. Signatures of Electrical Stimulation Driven Network Interactions in the Human Limbic System. J. Neurosci. 43, 6697–6711 (2023).

12. Huang, H. et al. Electrical Stimulation of Temporal and Limbic Circuitry Produces Distinct Responses in Human Ventral Temporal Cortex. J. Neurosci. 43, 4434–4447 (2023).

13. Trebaul, L. et al. Stimulation artifact correction method for estimation of early cortico-cortical evoked potentials. J. Neurosci. Methods 264, 94–102 (2016).

14. Prime, D., Woolfe, M., O’Keefe, S., Rowlands, D. & Dionisio, S. Quantifying volume conducted potential using stimulation artefact in cortico-cortical evoked potentials. J. Neurosci. Methods 337, 108639 (2020).

15. Jensen, M. A., Huang, H., Gregg, N. M., Muller, K. R., Hermes, D., & Miller, K. J. Approximating Signal Sources in Stereo-EEG Single Pulse Electrical Stimulation using Re-referencing and Spectral Analysis. bioRxivorg (2025): 2025–03.,.

16. Miller, K. J. et al. Canonical Response Parameterization: Quantifying the structure of responses to single-pulse intracranial electrical brain stimulation. PLoS Comput. Biol. 19, e1011105 (2023).

