## Supplemental Material for "The Human Insula Encodes Somatotopic Representation of Motor Execution with an Effector-Specific Connectomic Map to Primary Motor Cortex"

---

#### SUPPLEMENTAL METHODS

##### Colored somatotopic tuning

Somatotopic delineation ('tuning') was calculated for each channel in the following fashion: The individual $r^2$ values for each movement type were multiplied by $e^{i\pi/6}$ (hand), $e^{i\pi 5/6}$ (tongue) and $e^{i\pi 3/2}$ (foot) and added together. The magnitude of the resulting complex number defines the strength of somatotopic selectivity, and the phase angle of the complex number points to the movement (or pair of movements) that the channel is somatotopically specific for. This is illustrated in **Supplemental Figure 3**. Any negative $r^2$ values (i.e. broadband decreases with movement) were set to zero before calculation. Channels located in the precentral gyrus (on the gyral convexity or sulcal depth) with significant somatotopic selectivity were designated as somatotopically-tuned M1 sites and will be referred to as such henceforth.

##### Clinical electrical stimulation mapping

Three patients (s2, s3, s15) underwent extraoperative mapping with electrical stimulation as part of clinical workup. Interestingly, when electrical stimulation (50 Hz, 200 ms pulse duration, 2–5 second train, 1–5 mA) was delivered to insular contacts, no clear motor symptoms (e.g., muscle jerks, tremor, or negative motor phenomena) were elicited. Instead, patients reported sensory or vestibular sensations ("weird stomach sensation," "tingling throat sensation," "metallic taste," dizziness, vertigo). One patient (s15) exhibited delayed picture naming at higher stimulation intensities (2–3 mA) in the anterior insula. Comprehensive descriptions of these observations are provided in **Supplemental Table 3**.

##### Seizure onset zone and post-implantation management

Following the intracranial monitoring phase and a multidisciplinary review, the seizure onset zone was localized to a unilateral mesial temporal focus in seven patients, multifocal/generalized in five, combined mesial and neocortical temporal in three, and insular-frontal/temporal in two; one patient each was found to have pericentral, frontal, or mesial temporal onset.

#### SUPPLEMENTAL TABLES

**Suppl. Table 1.** Summary of patient demographics, seizure workup and implantation details.

| Patient | Age (y/o) | Sex | Dom. Hem. | Total lead n. | Insular lead* side (n) | MRI findings (3T) | SOZ hypothesis |
|---|---|---|---|---|---|---|---|
| s1 | 12 | M | L | 15 | R (2) | Bilateral paramedian midline ulegyria | multifocal (more posterior) |
| s2 | 18 | M | R | 12 | R (3) | s/p R ATL & stable tectal plate glioma | multifocal (mostly post temporal & insular) |
| s3 | 17 | F | L | 13 | R (2) | s/p R T lobectomy for PLNTY | R mesial temporal & insular |
| s4 | 12 | F | R | 17 | L (2) | L MTS & AG malformation | L mesial temporal |
| s5 | 13 | M | R | 16 | L (2) | L perinatal MCA infarct | Multifocal, frontal & mesial temporal |
| s6 | 17 | F | L | 16 | R (2) & L (2) | Nonlesional | R temporal |
| s7 | 20 | M | L | 12 | R (1) | s/p R PT lobectomy | R ant & post temporal |
| s8 | 15 | F | L | 12 | R (2) | R IF cavernoma | R frontal & mesial temporal |
| s9 | 17 | F | L | 15 | L (2) | L MTS | L mesial temporal |
| s10 | 13 | M | L | 16 | R (2) & L (2) | Nonlesional | generalized, mostly R mesial frontal |
| s11 | 17 | F | L | 13 | L (2) | Nonlesional | L temporal |
| s12 | 23 | M | L | 15 | R(3) | R hippocampal swelling; L hippocampal atrophy | L mesial temporal |
| s13 | 37 | M | L | 15 | R (2) | Nonlesional | R mesial temporal |
| s14 | 11 | M | L | 12 | L (2) | Nonlesional | L pericentral |
| s15 | 15 | F | L | 10 | L (2) | L MTS | L mesial temporal |
| s16 | 22 | F | L | 14 | R (2) | s/p shunt at young age for hydrocephalus | R mesial temporal |
| s17 | 18 | F | L | 15 | L (2) | Bifrontal WM gliosis & atrophy | multifocal |

**Abbreviations:** AG: angular gyrus; ATL: anterior temporal lobectomy; GWM: gray-white matter; IF: inferior frontal; L: left; MTS: mesial temporal sclerosis; PLNTY: Polymorphous low-grade neuroepithelial tumor of the young; PT: parietal; R: right; SF: superior frontal; SOZ: seizure onset zone; STG: superior temporal gyrus; T: temporal; WM: white matter

*All leads were made from DIXI Medical, France, except for s16 which were made from PMT Corporation, USA.

**Suppl. Table 2.** Summary of identified somatotopically-tuned sites in the insula and primary motor cortex across patients.

| Patient | Primary motor cortex (M1) | | | Insular cortex | | | |
|---|---|---|---|---|---|---|---|
| | Hand | Tongue | Foot | Hand | Tongue | Foot | Inter-effector* |
| s1 | Yes | No | No | Yes | Yes | No | No |
| s2** | Yes | No | No | Yes | Yes | Yes | Yes |
| s3 | Yes | Yes | No | Yes | Yes | No | Yes |
| s4 | Yes | Yes | No | Yes | Yes | No | No |
| s5 | Yes | Yes | No | Yes | Yes | No | No |
| s6 | No | No | No | Yes | Yes | No | No |
| s7 | Yes | Yes | No | No | Yes | No | No |
| s8 | Yes | No | No | Yes | Yes | No | No |
| s9 | Yes | No | No | No† | No† | No | No |
| s10 | Yes | No | No | No | Yes | Yes | No |
| s11 | No | No | No | Yes | Yes | No | No |
| s12 | Yes | Yes | No | Yes | Yes | No | No |
| s13 | No | Yes | No | No | Yes | No | No |
| s14 | Yes | Yes | No | Yes | Yes | No | No |
| s15 | Yes | Yes | No | No | Yes | No | Yes |
| s16 | No | Yes | No | Yes | Yes | Yes | No |
| s17 | No | No | No | No | No | No | Yes |

*Defined as $r^2$ value > 0.10 and p-value < 0.05 for hand & tongue & foot movement.
** s2, s8, s9,  did not have EMG of sufficient quality to allow for brain-movement latency estimation
†Significant hand and tongue channels overlapped.

**Suppl. Table 3.** Summary of reaction times (cue to movement onset interval) for each patient

| Patient | Reaction time in ms, mean (st. deviation) | | |
|---|---|---|---|
| | **Hand** | **Tongue** | **Foot** |
| s1 | 725 (420) | 726 (415) | 778 (558) |
| s2 | 415 (138) | Suboptimal EMG | 371 (127) |
| s3 | 393 (131) | 505 (254) | 419 (84) |
| s4 | 744 (442) | Suboptimal EMG | 670 (225) |
| s5 | 413 (168) | 473 (95) | 592 (360) |
| s6 | 395 (153) | 316 (79) | 415 (179) |
| s7 | 477 (122) | 411 (97) | 494 (136) |
| s8 | No EMG | | |
| s9 | 402 (249) | Suboptimal EMG | 527 (495) |
| s10 | 641 (331) | 490 (144) | 608 (499) |
| s11 | 287 (91) | 215 (377) | 429 (138) |
| s12 | 687 (131) | 564 (511) | 588 (128) |
| s13 | 268 (66) | 360 (101) | 317 (93) |
| s14 | 731 (216) | 715 (210) | 710 (195) |
| s15 | 444 (133) | 504 (205) | 372 (112) |
| s16 | 590 (224) | 677 (330) | 499 (151) |
| s17 | 721 (256) | 593 (177) | 681 (208) |

**Suppl. Table 4.** Summary of clinical electrical stimulation mapping results in three subjects and comparison with findings from our behavioral motor task.

| Subject | Stimulated Insular lead | Evoked response | Motor task results |
|---|---|---|---|
| s2 | RV1-2 | Dizziness, vertigo | Not significant |
| | RV3-4 | Dizziness, vertigo | Not significant |
| | RV5-6 | Dizziness, vertigo | Not significant |
| | RMI1-2 | Metallic taste | Significant hand activation |
| | RMI3-4 | Metallic taste | Significant hand & tongue activation |
| | RMI5-6 | none | Significant tongue activation |
| | RMI7-8 | none | Significant tongue activation |
| | RP1-2 | Stomach sensation, metallic taste | Significant hand activation |
| | RP3-4 | Stomach sensation, metallic taste | Not significant |
| | RP5-6 | Stomach sensation, metallic taste | Not significant |
| s3 | RV1-2 | none | Significant hand activation |
| | RV3-4 | none | Significant tongue activation |
| | RV5-6 | none | Not significant |
| | RP1-2 | none | Cross-modal site |
| | RP3-4 | none | Significant hand & tongue activation |
| | RP5-6 | Tingling throat sensation | Significant tongue activation |
| s15 | LV1-2 | Slight slower reaction with naming | Not significant |
| | LV3-4 | Slower naming response, dizzy sensation | Significant hand & tongue activation |
| | LP1-2 | ‘fuzzy’ sensation | Not significant |
| | LP3-4 | ‘fuzzy’ sensation | Significant hand activation |

#### SUPPLEMENTAL FIGURES

#### FIGURE S1

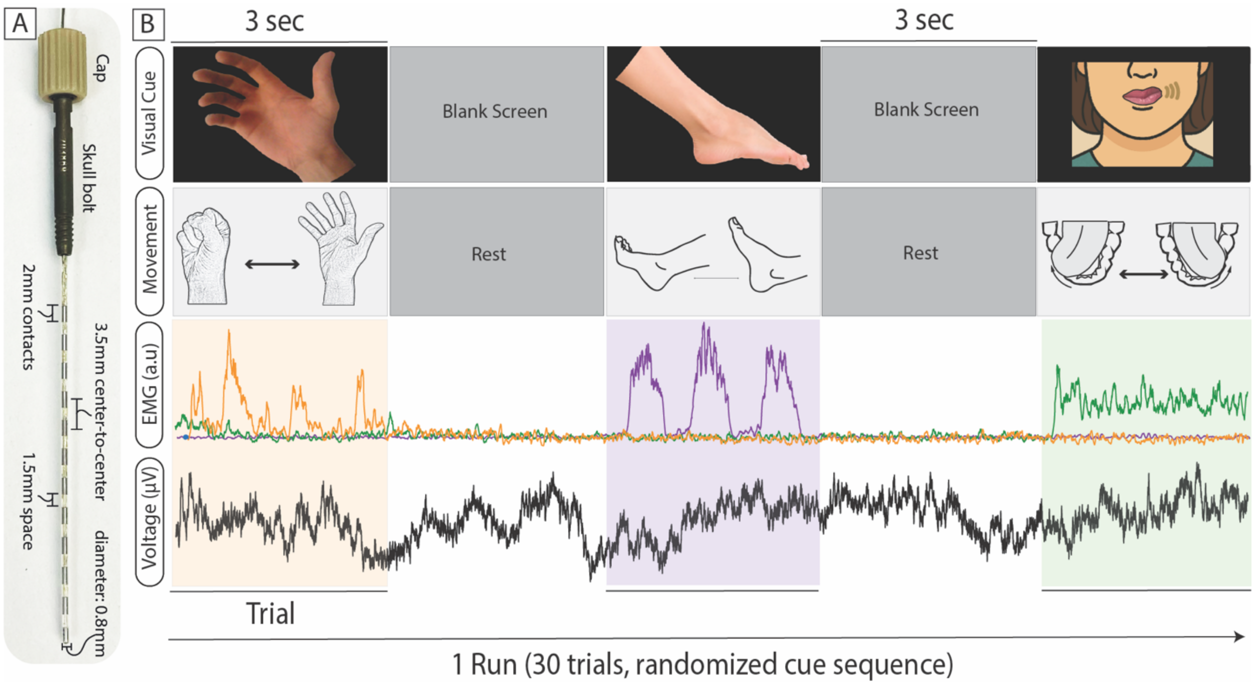


**Suppl. Figure 1.** (A) Macroscopic view and specifications of the Dixi stereo-EEG lead which was implanted in our cohort. (B) Overview of the motor task design. An experimental run consisted of thirty, randomly shuffled movement-rest trial periods (10 per hand, tongue and foot). Movement of each body part was captured with synchronized EMG, which was the basis of segmenting our data into movement and rest epochs. Each subject completed two experimental runs.

#### FIGURE S2

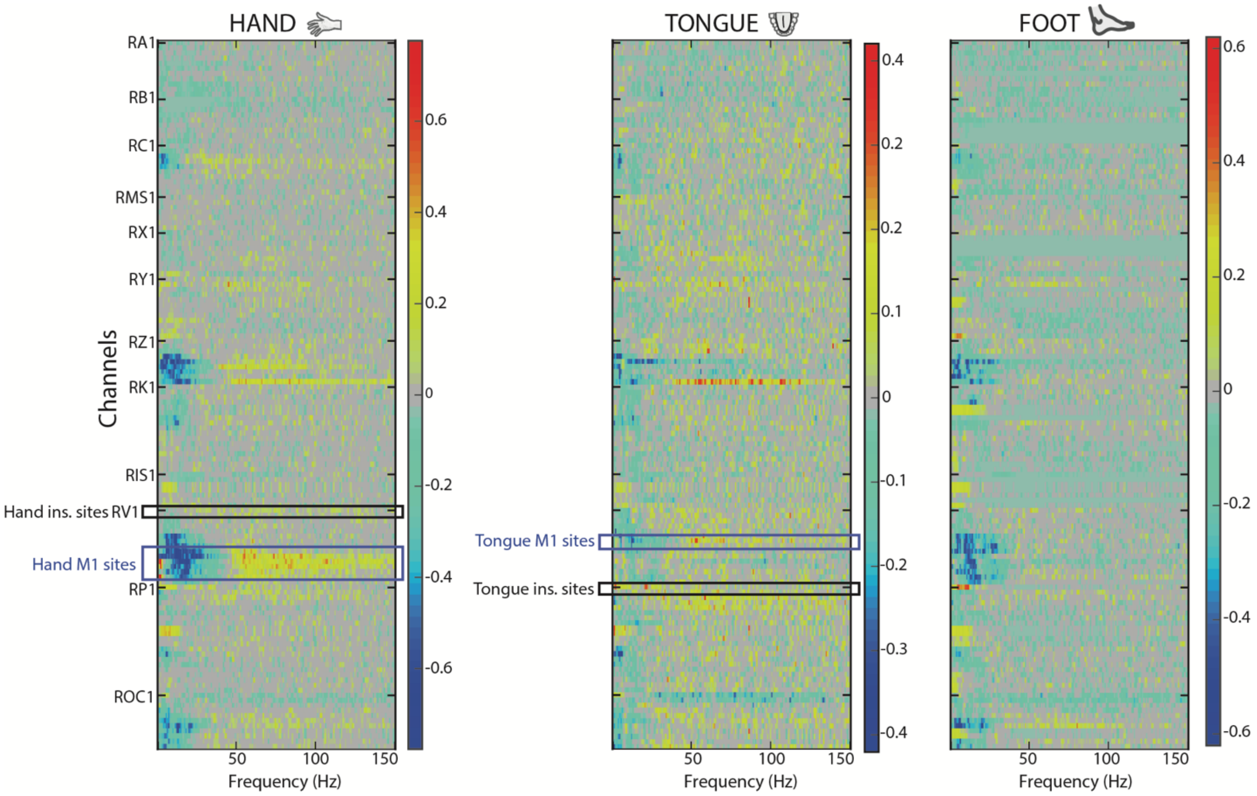


**Suppl. Figure 2.** $R^2$ activation feature maps (s1) demonstrating the power change in 1Hz frequency bins per channel, for each movement type. It is noted that there is significant decrease in low frequency oscillations and increase in broadband high-frequency range in the primary motor cortical sites during hand and tongue movement compared to rest. We did not observe a shift in oscillations with movement in the corresponding insular sites.

**FIGURE S3**

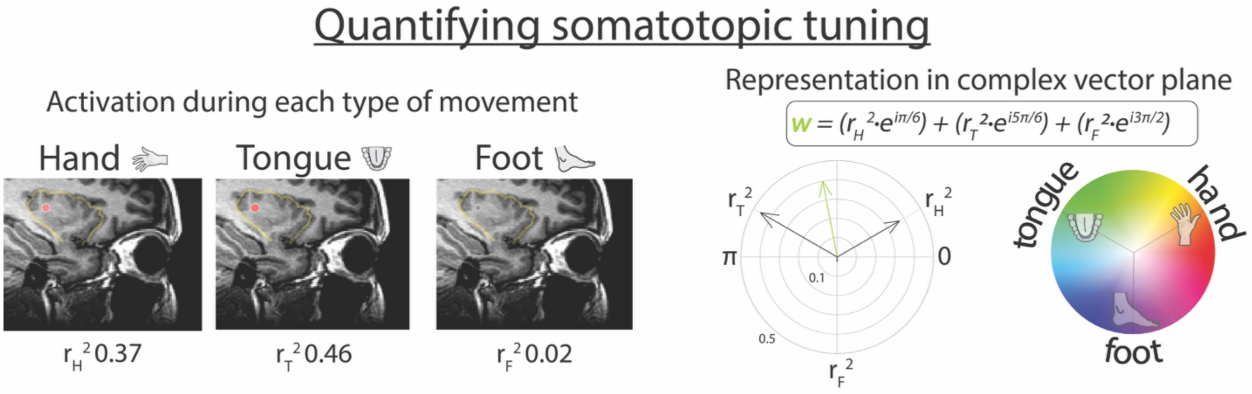


**Suppl. Figure 3.** Somatotopic tuning at each channel can be quantified by representing the $r^2$ values associated with each movement type as vectors in the complex plane. For each vector, its magnitude is equal to the $r^2$ value and its phase is assigned based on the movement type. The linear summation of the hand, tongue and foot vectors produces a new vector (green arrow). The magnitude of the resulting vector defines the strength of somatotopic selectivity, and the phase angle points to the movement that the channel is somatotopically specific for.

#### FIGURE S4

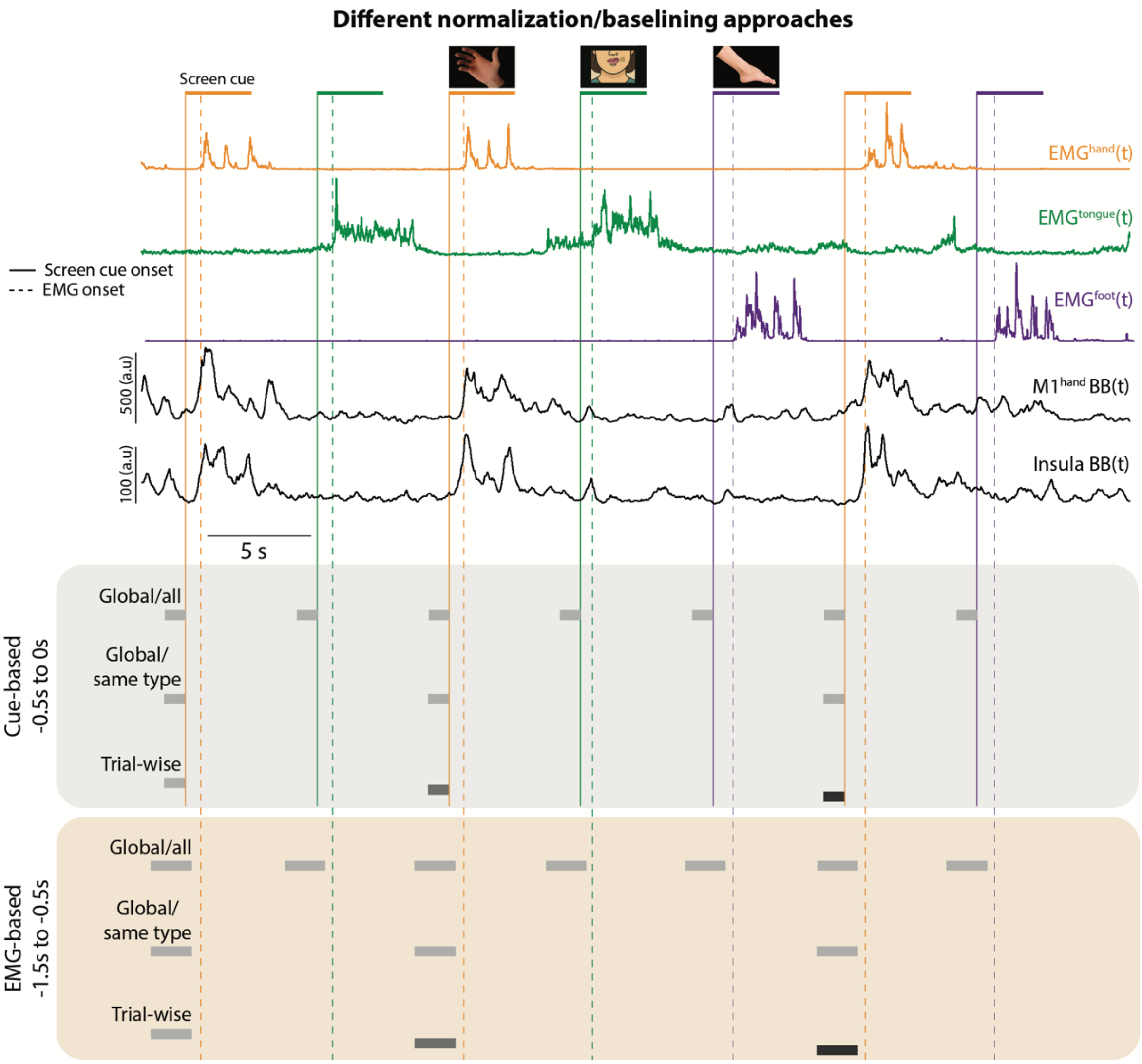


**Suppl. Figure 4.** Different normalization/baselining approaches exist for neural recordings depending on the paradigm and research question. They can be broadly divided into cue-based and behavioral onset-based. These can be further subdivided into a) global and considering all trials, b) global and considering only trials of the same type (e.g. hand movement) and c) trial-wise, where each trial is normalized to each own baseline. In the present analysis, we used global-all, cue-based normalization. BB: broadband power. EMG: electromyogram.

#### FIGURE S5

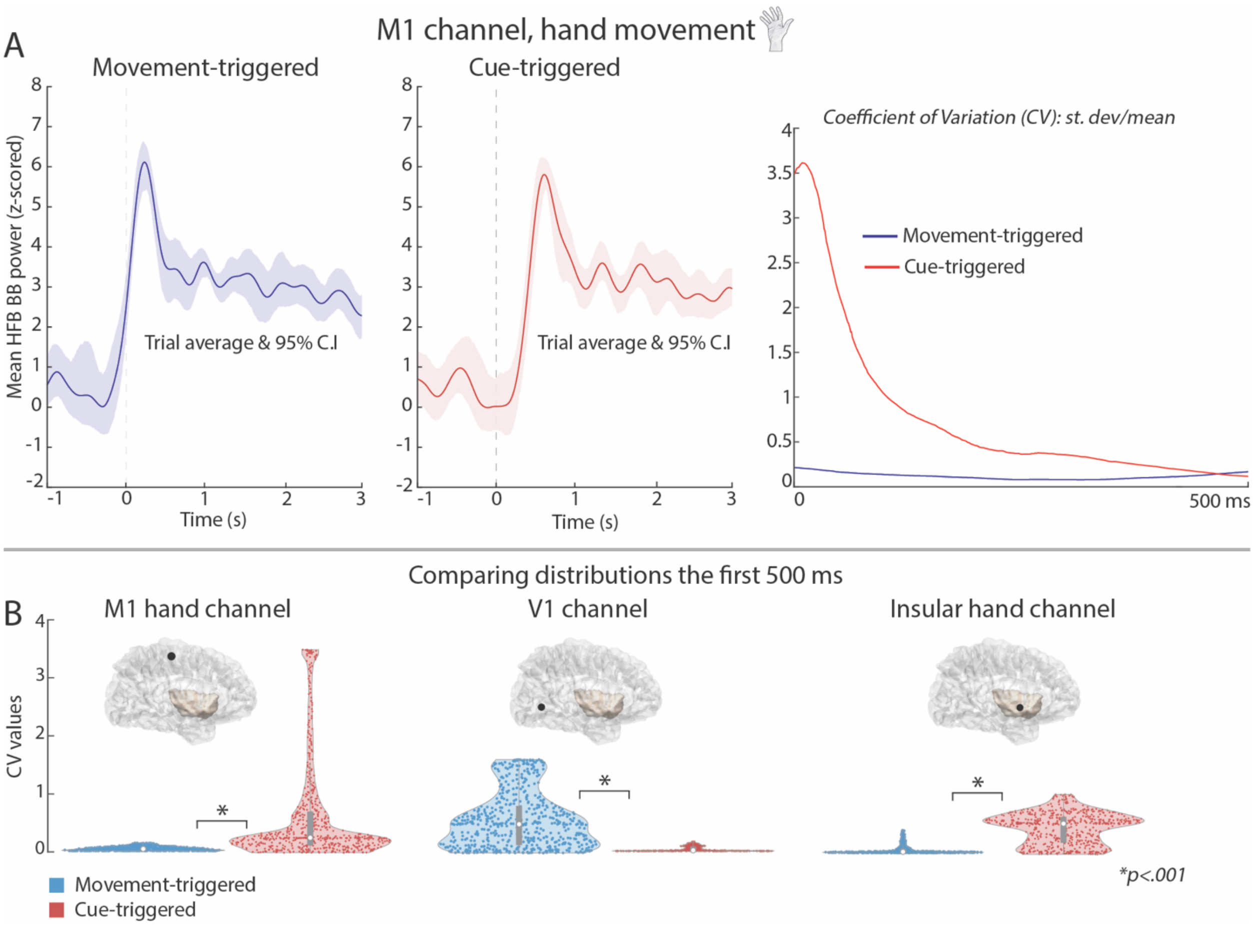


**Suppl. Figure 5.** Comparison of movement-triggered versus screen cue (stim)-triggered average traces along with 95% confidence intervals. (A) The coefficient of variation during the first 500 ms following each type of trigger is computed and the two distributions are compared using bootstrap resampling statistics. (B) The M1 and insular hand channel activity showed significantly lower spread with movement triggering, which means they are movement- and not stim- locked, while the V1 (primary visual cortex) channel activity, as expected, was stim-locked.

#### FIGURE S6

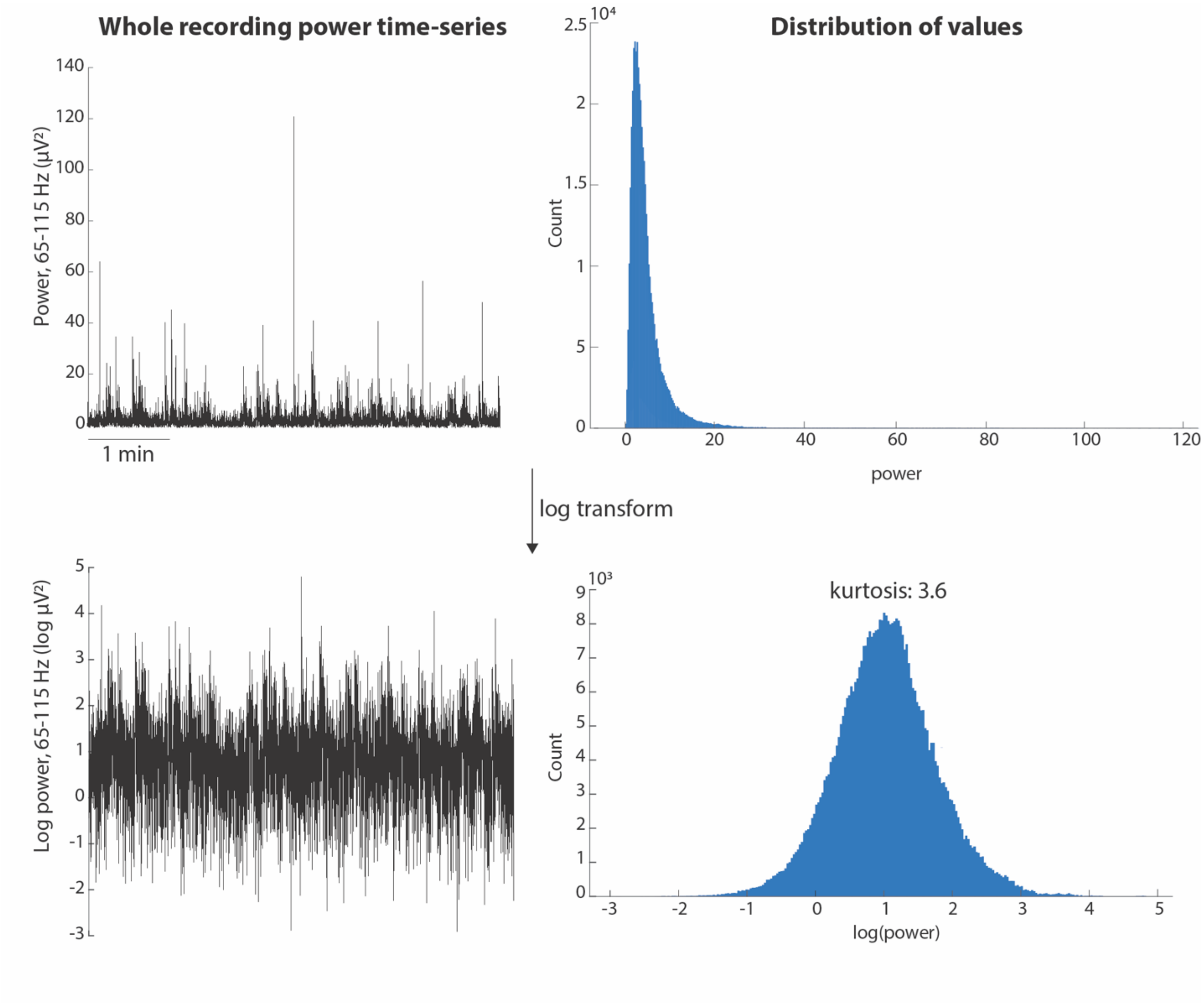


**Suppl. Figure 6.** The high-frequency broadband power time series during the behavioral task (subject 3) shows a skewed distribution (hand M1 channel in this example). Following log transformation, the values are normally distributed.

#### FIGURE S7

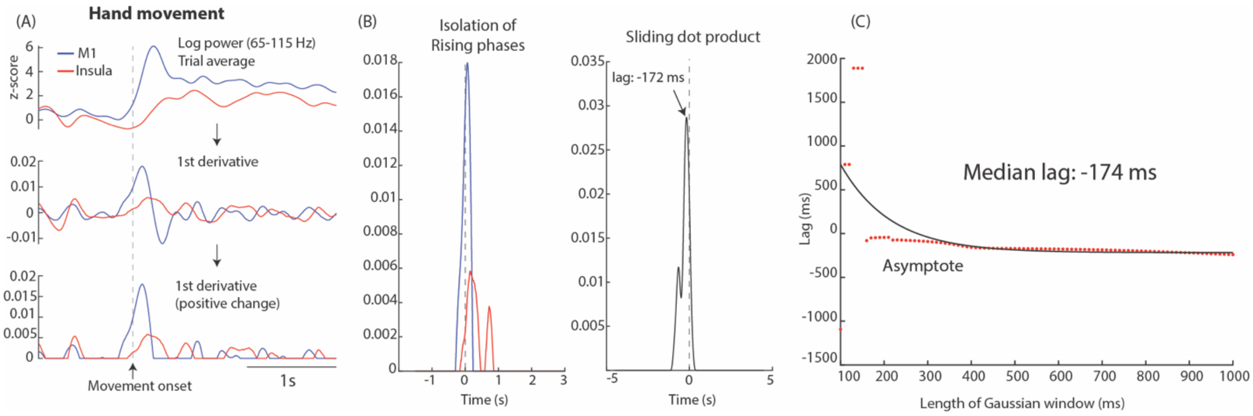


**Suppl. Figure 7.** Demonstration of the 'rising phases isolation' method. (A) The first derivative is computed for the movement-triggered average traces for somatotopic M1 and insular channels. Negative values are set to zero in order to identify periods of positive (rising) change. (B) The largest segment that also contains the global maximum within 1 second after movement onset was isolated for each channel and the sliding dot product was computed. The time point of maximum correlation corresponds to the lag between M1 and insular activation for a specific movement type. (C) The process was repeated using different lengths of Gaussian kernels (100-1000 ms) to smoothen the power time series and evaluate the sensitivity of our results. The median lag from this distribution approximated the value obtained using the 500 ms smoothing window used in our analysis.

**FIGURE S8**

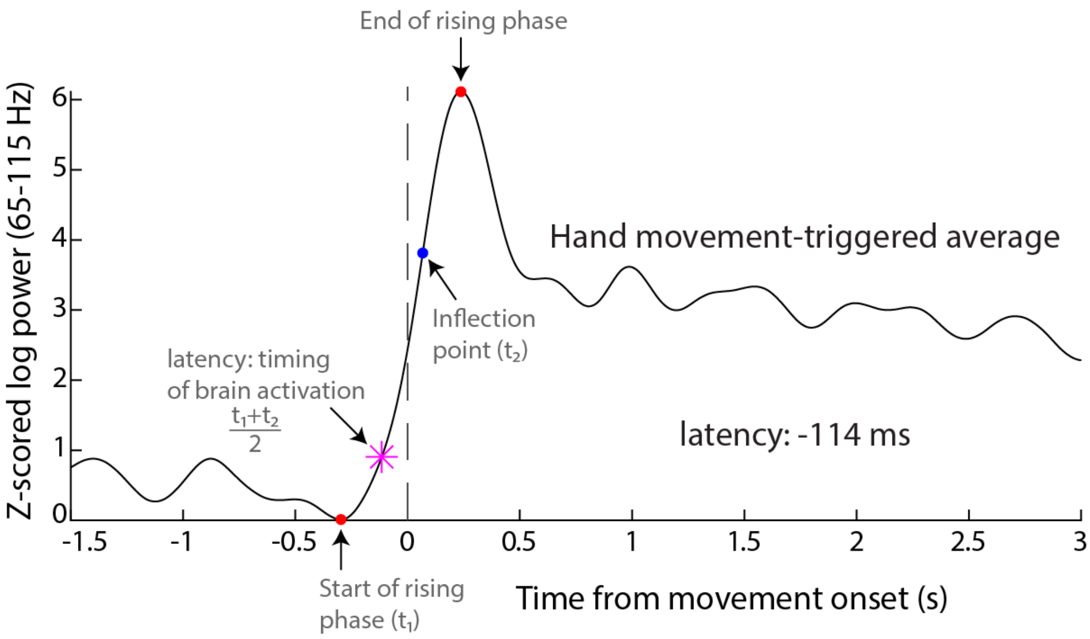


**Suppl. Figure 8.** Demonstration of the method for calculating for timing of brain activation (latency) with respect to movement onset. To calculate the latency (pink asterisk), we identified two key timepoints in the rising phase of the movement triggered average trace: the onset (red circle, t1) and the inflection point where the second derivative equals zero (blue circle, t2). The latency was defined as the midpoint between these two timepoints. In this example, brain activity in the M1 hand channel preceded movement onset by 114 ms.

**FIGURE S9**

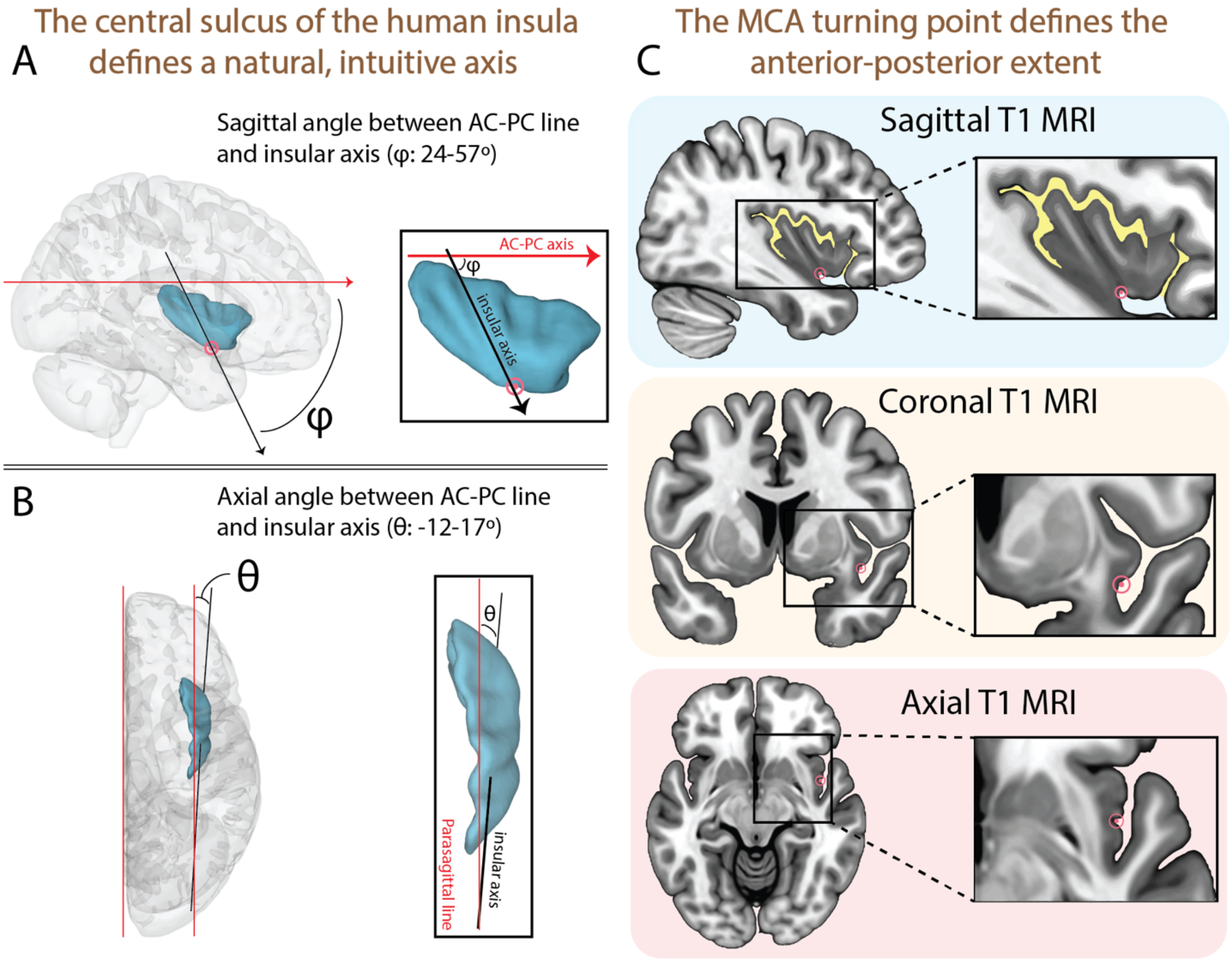


**Suppl. Figure 9.** Angles defined by the insular axis. The insular axis passing through the central insular sulcus forms (A) a sagittal angle φ and (B) an axial angle θ with the AC-PC line, which are used to rotate each individual brain from the AC-PC into the insular stereotactic space. (C) An indentation is typically observed in the inferior corner, where the middle cerebral artery (MCA) takes a steep turn before its bifurcation/trifurcation. This point was selected to define the anterior/posterior extent (i.e. y=0).

#### FIGURE S10

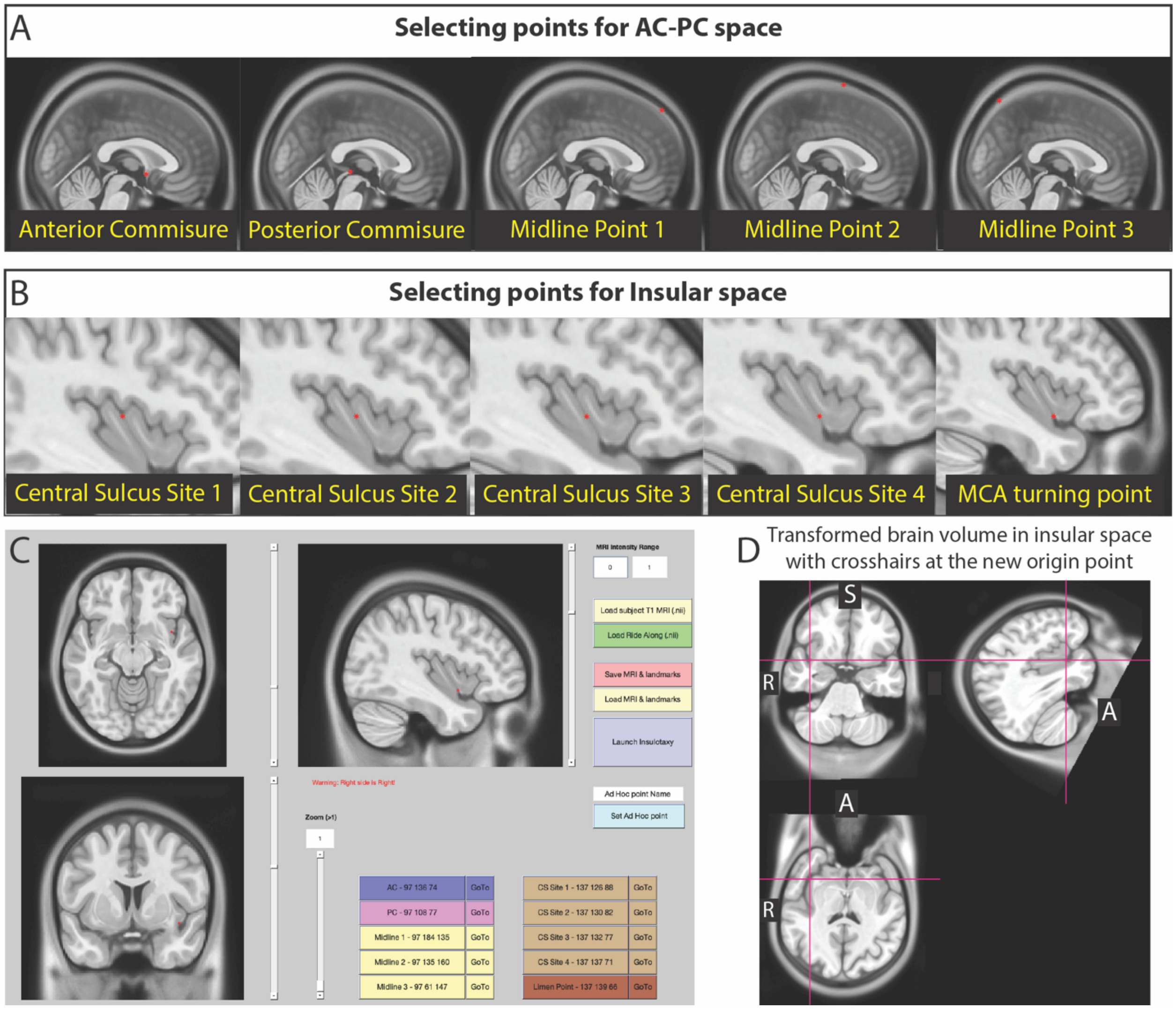


**Suppl. Figure 10.** Demonstration of the graphical user interface used to convert each individual brain volume into the insular stereotactic space based on an affine local transformation. (A) First, the user identifies five points necessary to rotate into the AC-PC stereotactic space, (B) followed by five points to rotate into the insular space. (C) In this example, we used the right insula in MNI-152 2009c, nonlinear asymmetric, skull-stripped brain. (D) Once transformation is complete, the brain volume is successfully rotated into the new coordinate space, which can be used to visualize group-level data.

#### FIGURE S11

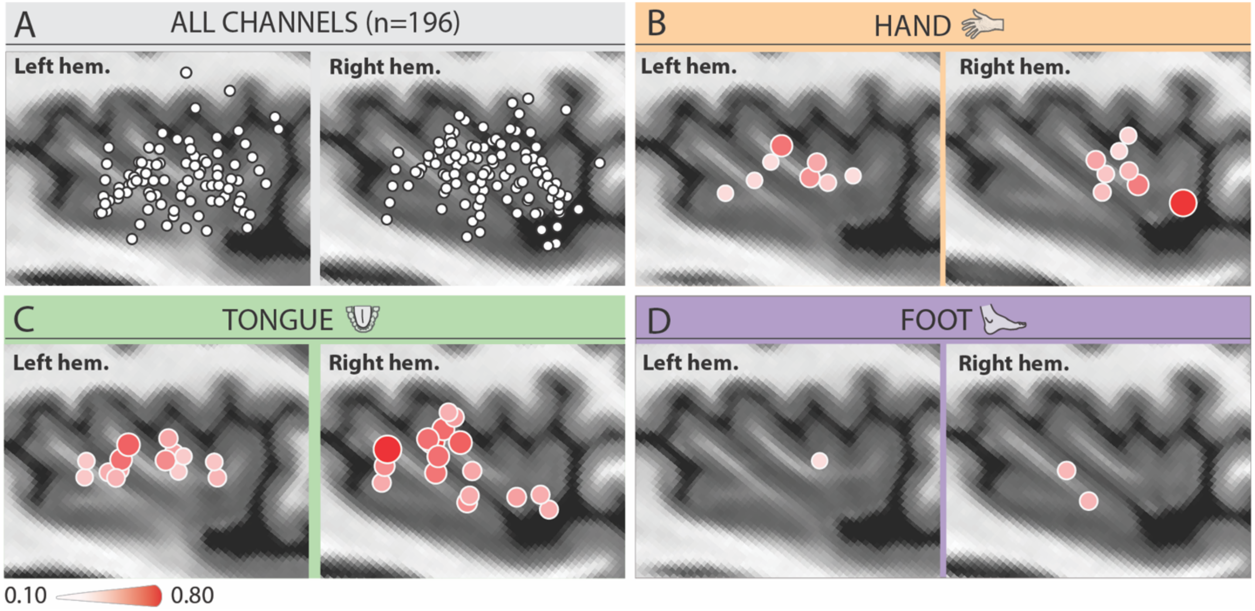


**Suppl. Figure 11.** (A) Positions of bipolar channels, (C) hand, (D) tongue and (E) foot $r^2$ activation maps for the left and right hemisphere, respectively.

#### FIGURE S12

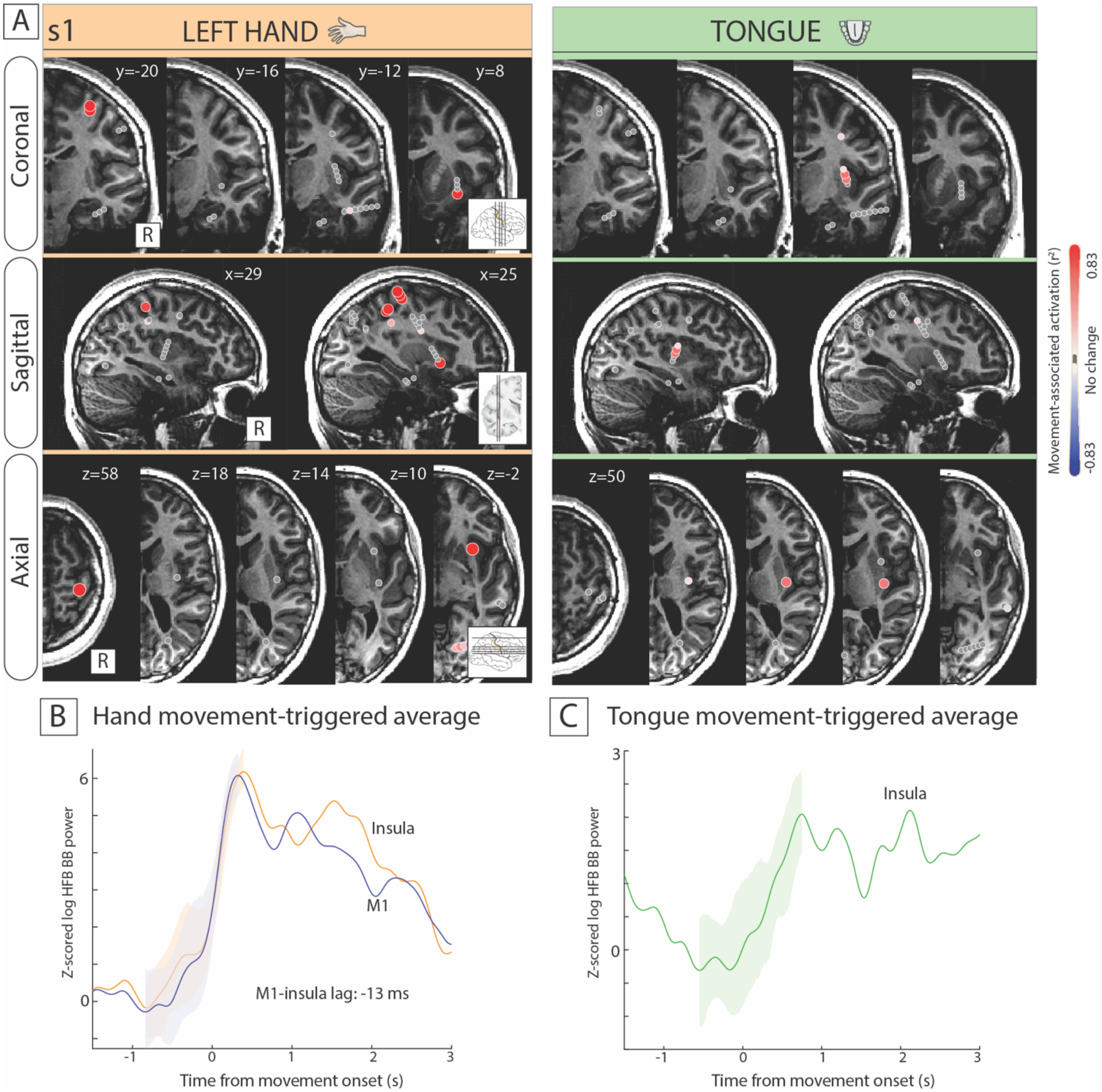


**Suppl. Figure 12.** Strong hand-tuned insular site with broadband activation preceding the corresponding hand M1 site (s1). (A) Activation $r^2$ maps for hand (orange) and tongue (green) movement in coronal, sagittal and axial planes (4 mm slices). (B) Movement-cue-triggered average (-1.5 s to 3 s) high-frequency broadband responses comparing the activity of the hand-tuned sites, along with the standard error of the mean during the rising phase. C) Movement-triggered average high-frequency broadband response for the tongue insular site.

#### FIGURE S13

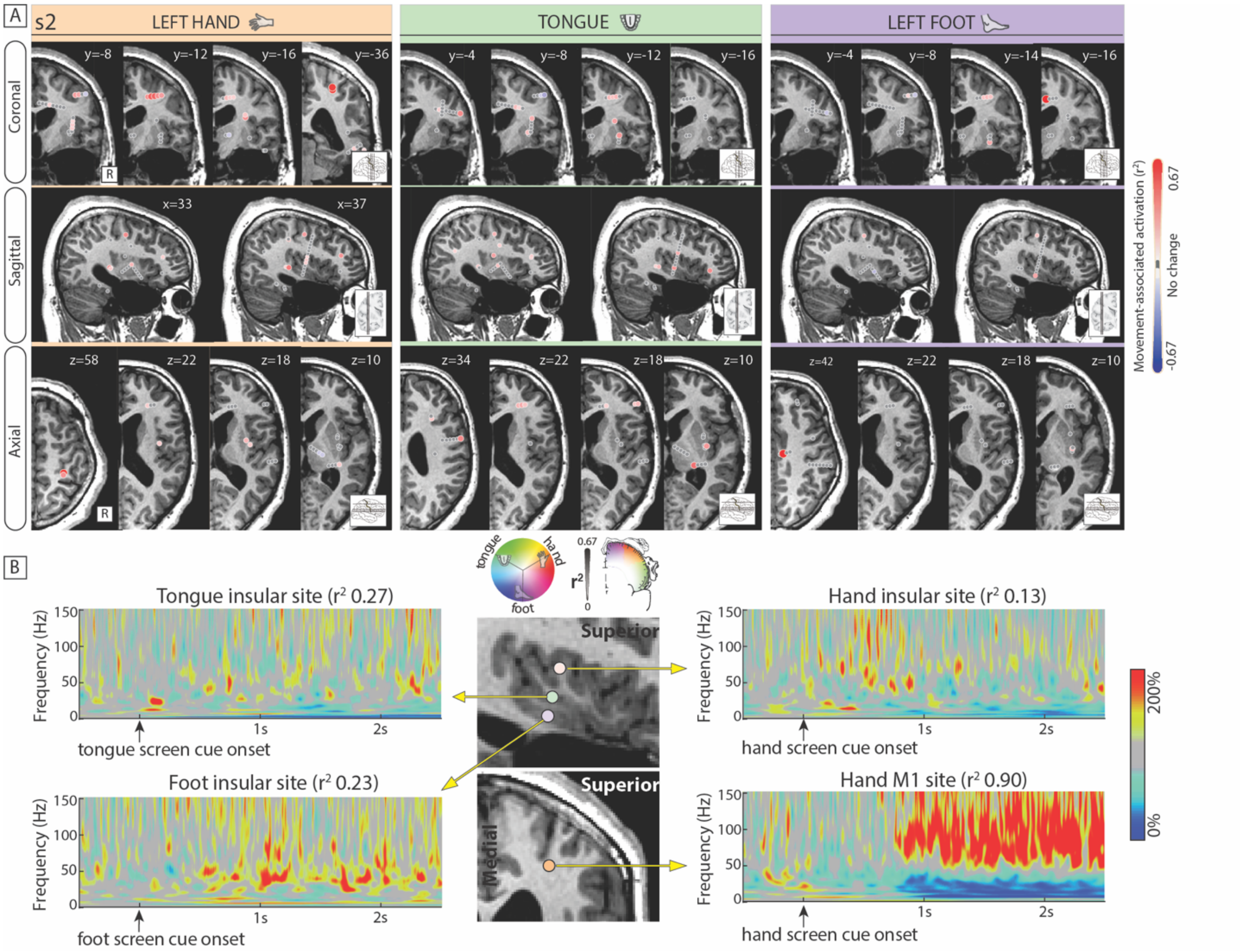


**Suppl. Figure 13.** Hand, tongue and foot representation in subject 2. (A) Activation $r^2$ maps for hand (orange) and tongue (green) movement in coronal, sagittal and axial planes (4 mm slices). The peri-insular sulcus is shown in yellow. (B) Screen cue-triggered average (-1.5 s to 1.5 s) spectrograms for the hand M1, hand insular, tongue insular and foot insular sites.

#### FIGURE S14

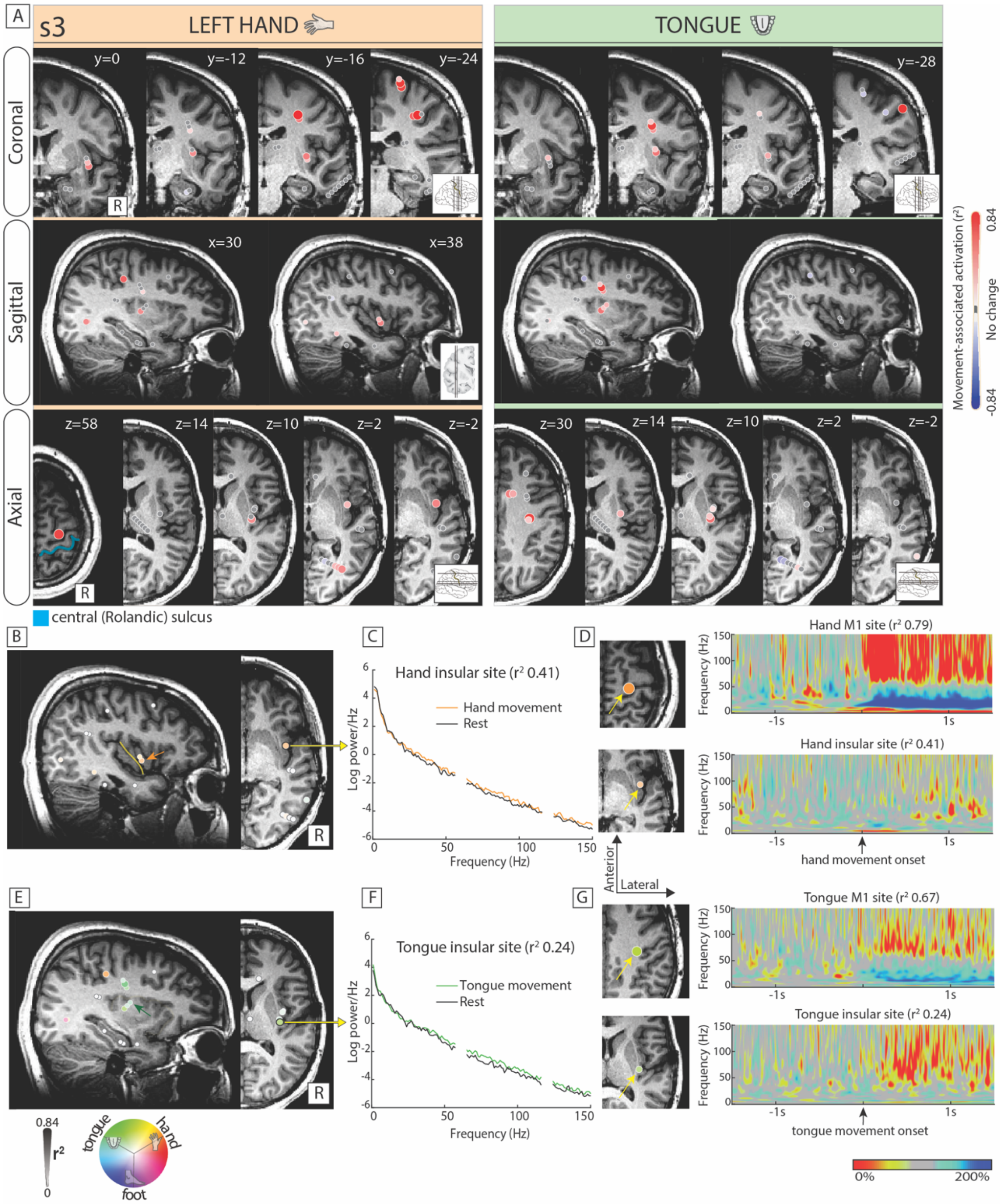


**Suppl Figure 14.** Hand and tongue motor representation in subject 3. (A) Activation $r^2$ maps for hand (orange) and tongue (green) and movement in coronal, sagittal and axial planes (4 mm slices). The central sulcus is shown in blue. (B) Hand-tuned sites (orange) are noted in the middle part of the insula, just anterior to its central sulcus (in yellow). (C) Trial-average power spectrum density of a hand-tuned channel. (D) Movement-triggered average (MTA; -1.5 s to 1.5 s) spectrograms during hand movement for the hand-tuned primary motor cortical and insular sites. (E-G) The same as in panels C-D, but for tongue-selective channels.

#### FIGURE S15

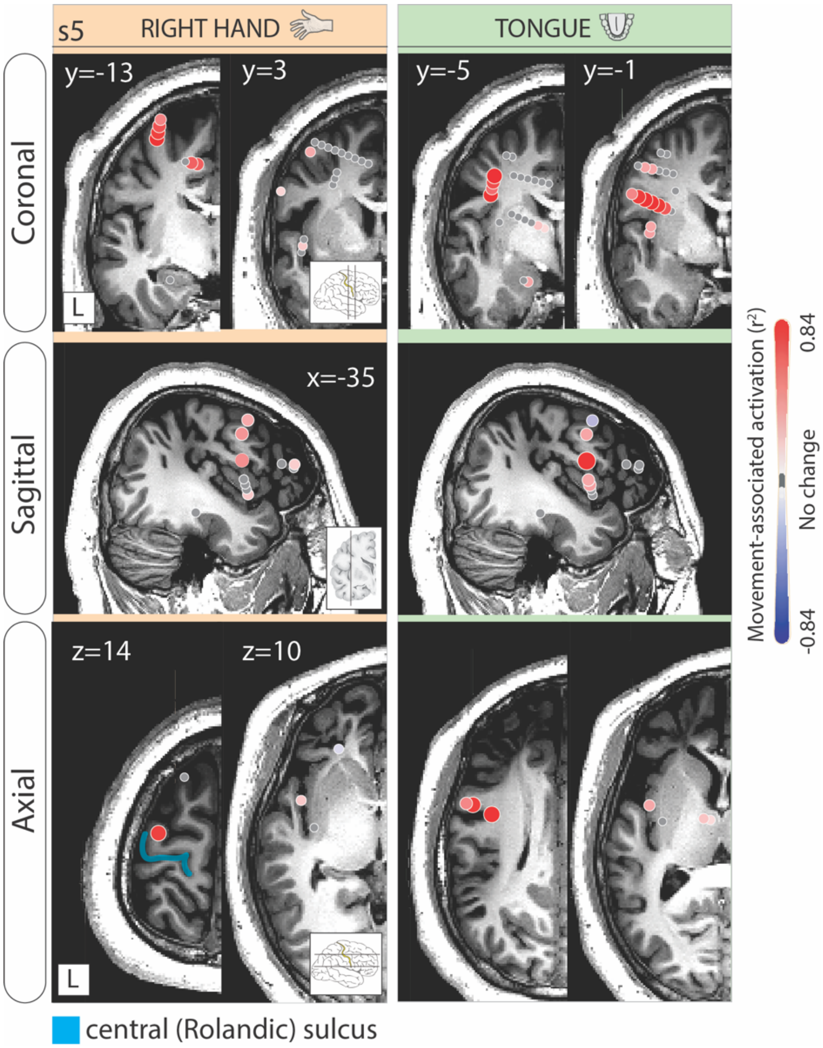


**Suppl Figure 15.** Insular activation with hand and tongue motor execution (s4). Activation $r^2$ maps for hand (orange) and tongue (green) movement in coronal, sagittal and axial planes (4 mm slices).

#### FIGURE S16

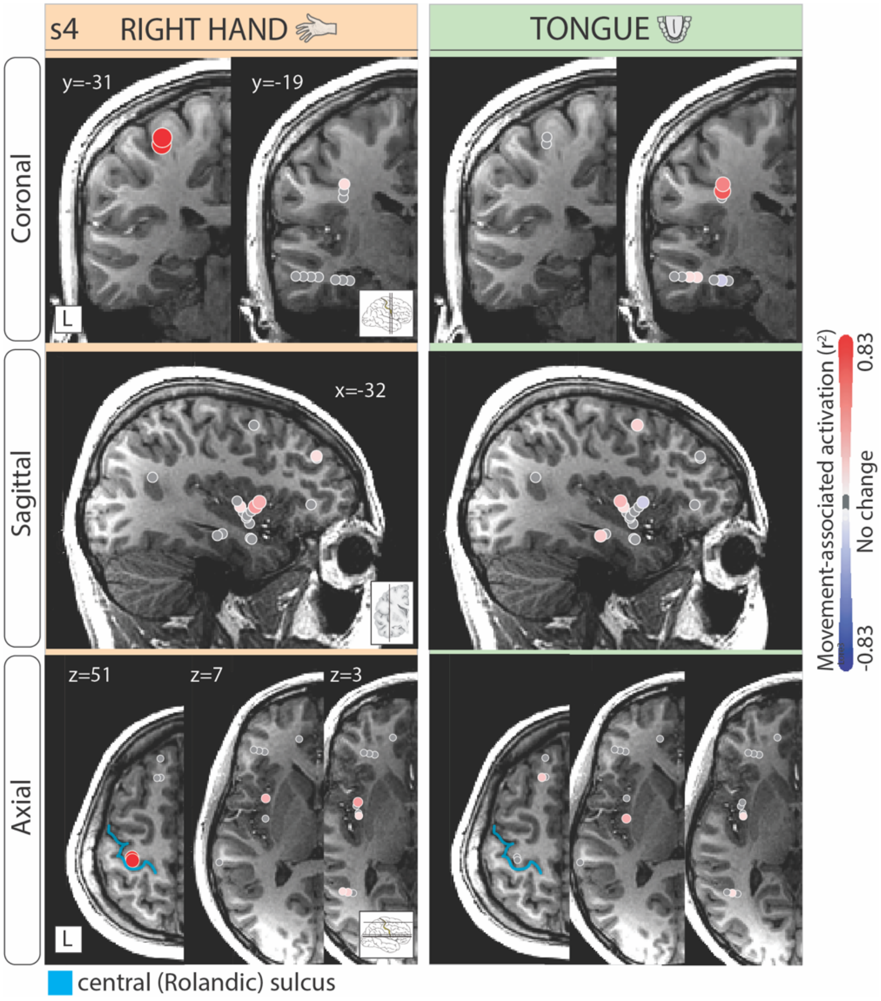


**Suppl Figure 16.** Insular activation with hand and tongue motor execution (s5). Activation $r^2$ maps for hand (orange) and tongue (green) movement in coronal, sagittal and axial planes (4 mm slices).

### FIGURE S17

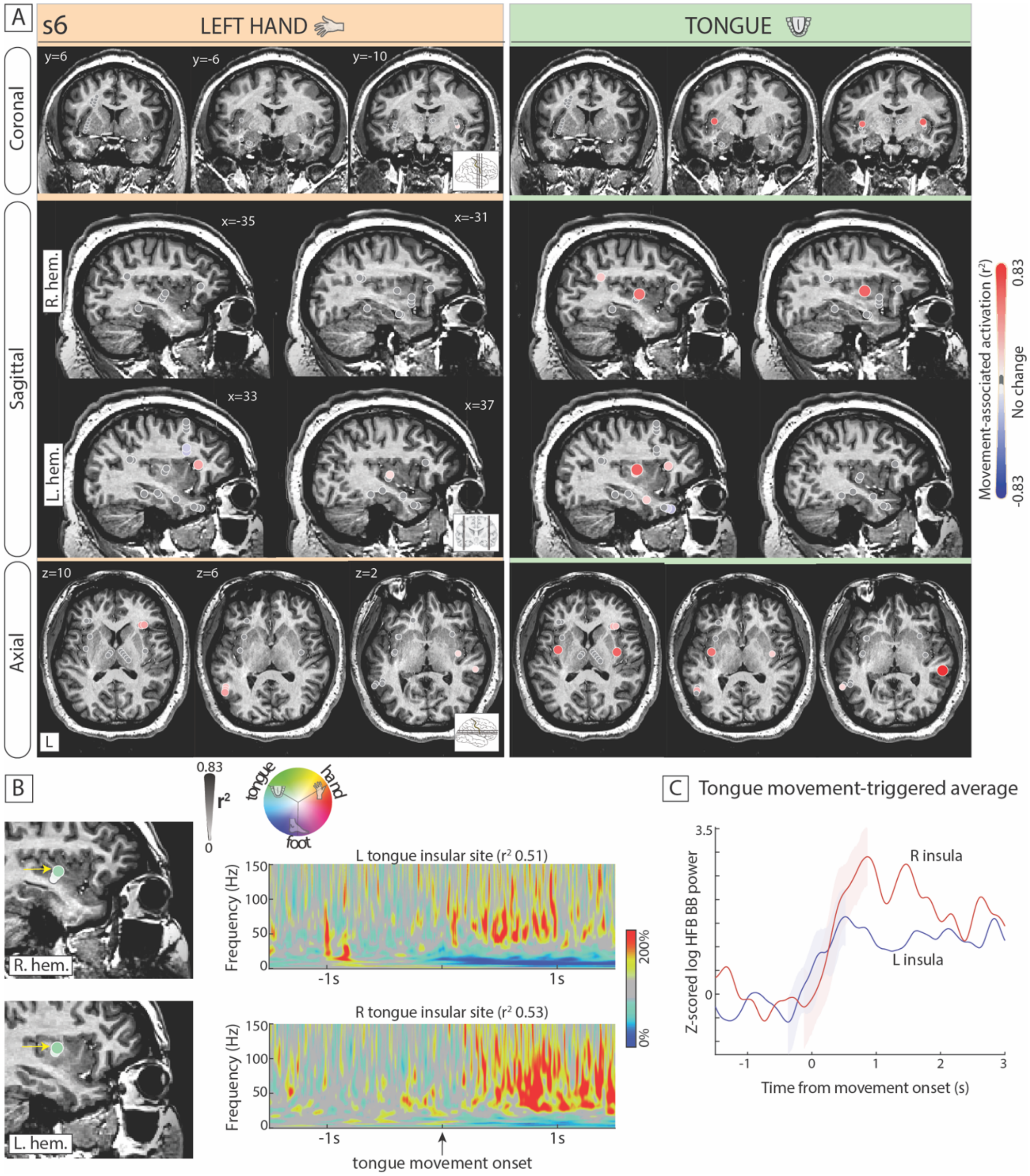


**Suppl Figure 17.** Insular motor activity is bilateral for tongue movement and contralateral for hand movement (s6). (A) Activation $r^2$ maps for hand (orange) and tongue (green) movement in coronal, sagittal and axial planes (4 mm slices). Channel activation was only contralateral to hand movement, while there was strong bilateral tongue activation. (B) Movement-triggered average (-1.5 s to 1.5 s) spectrograms during tongue movement for bilateral tongue-tuned insular sites. (C) Movement cue-triggered average (-0.5 s to 3s) high-frequency broadband responses, along with the standard error of the mean during the rising phase, comparing the activity of right vs left tongue-tuned insular sites.

#### FIGURE S18

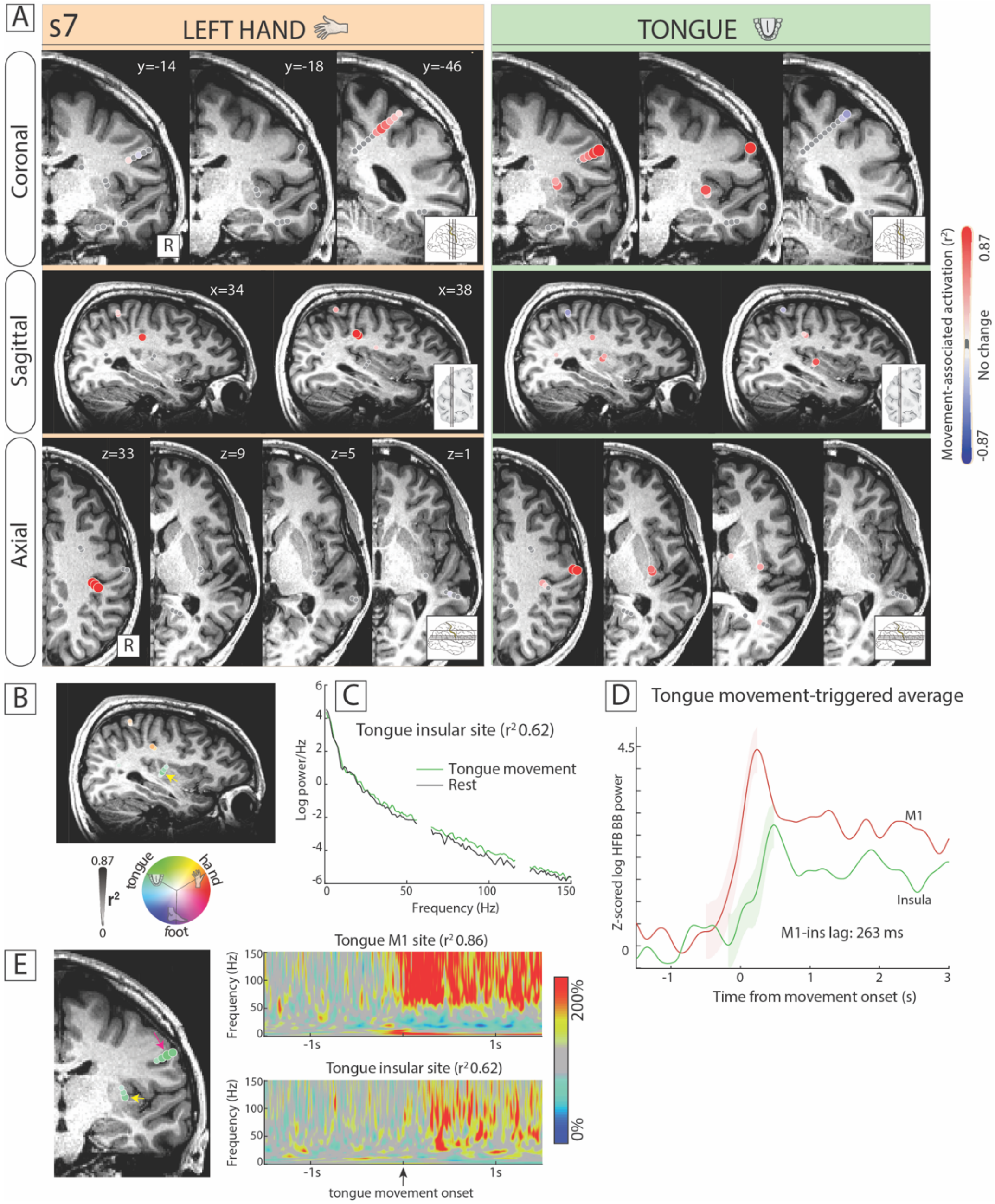


**Suppl. Figure 18.** Strong motor representation of the tongue in the posterior insula (s7). (A) Activation $r^2$ maps for hand (orange) and tongue (green) movement in coronal, sagittal and axial planes (4 mm slices). (B) Tongue-selective channels (yellow arrow) localized in the posterior insula. (C) Trial-average power spectrum density plot for tongue movement vs rest in the tongue insular site with the highest $r^2$ value. (D) Movement-triggered average (-0.5 s to 3 s) high-frequency broadband responses, along with the standard error of the mean during the rising phase, comparing the activity of the same insular site to M1 activity. (E) Movement-triggered average (-1.5 s to 1.5 s) spectrograms for the tongue M1 (pink arrow) and insular site (yellow arrow).

#### FIGURE S19

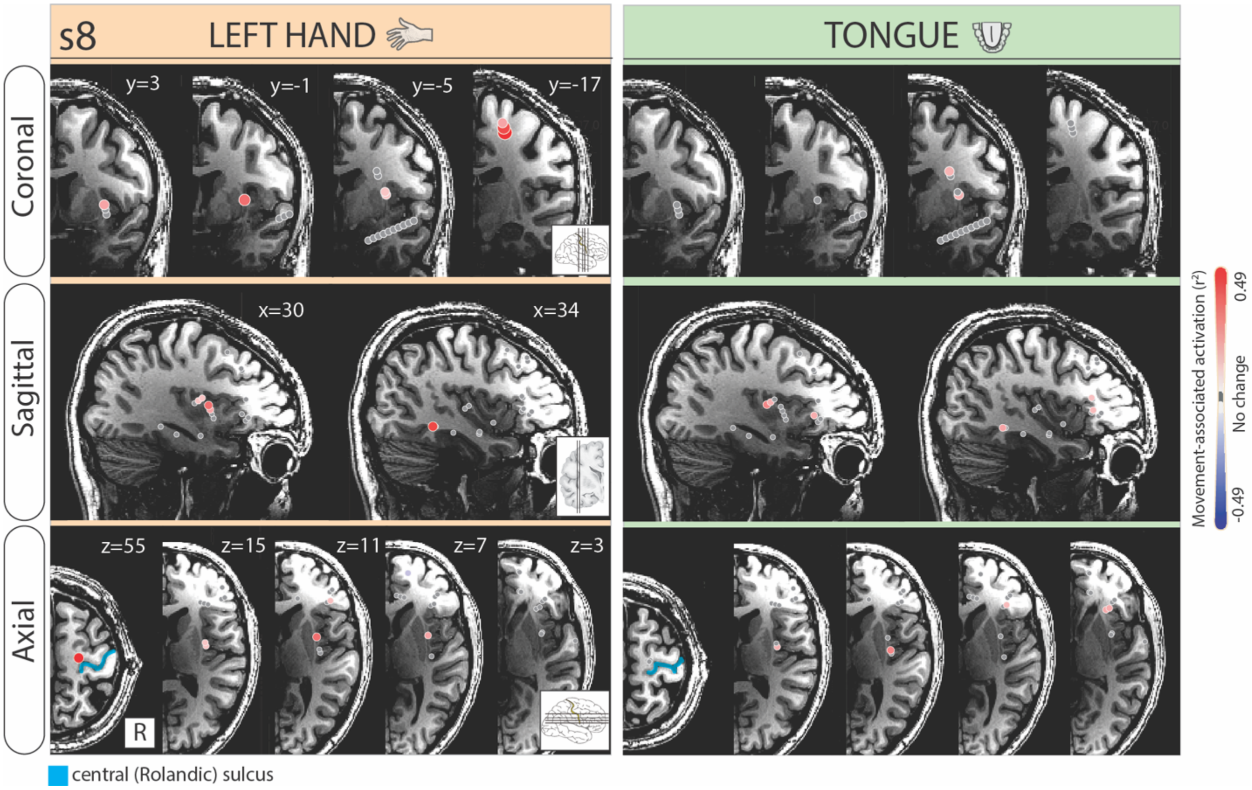


**Suppl. Figure 19.** Insular activation with hand and tongue motor execution (s8). Activation $r^2$ maps for hand (orange) and tongue (green) movement in coronal, sagittal and axial planes (4 mm slices).

#### FIGURE S20

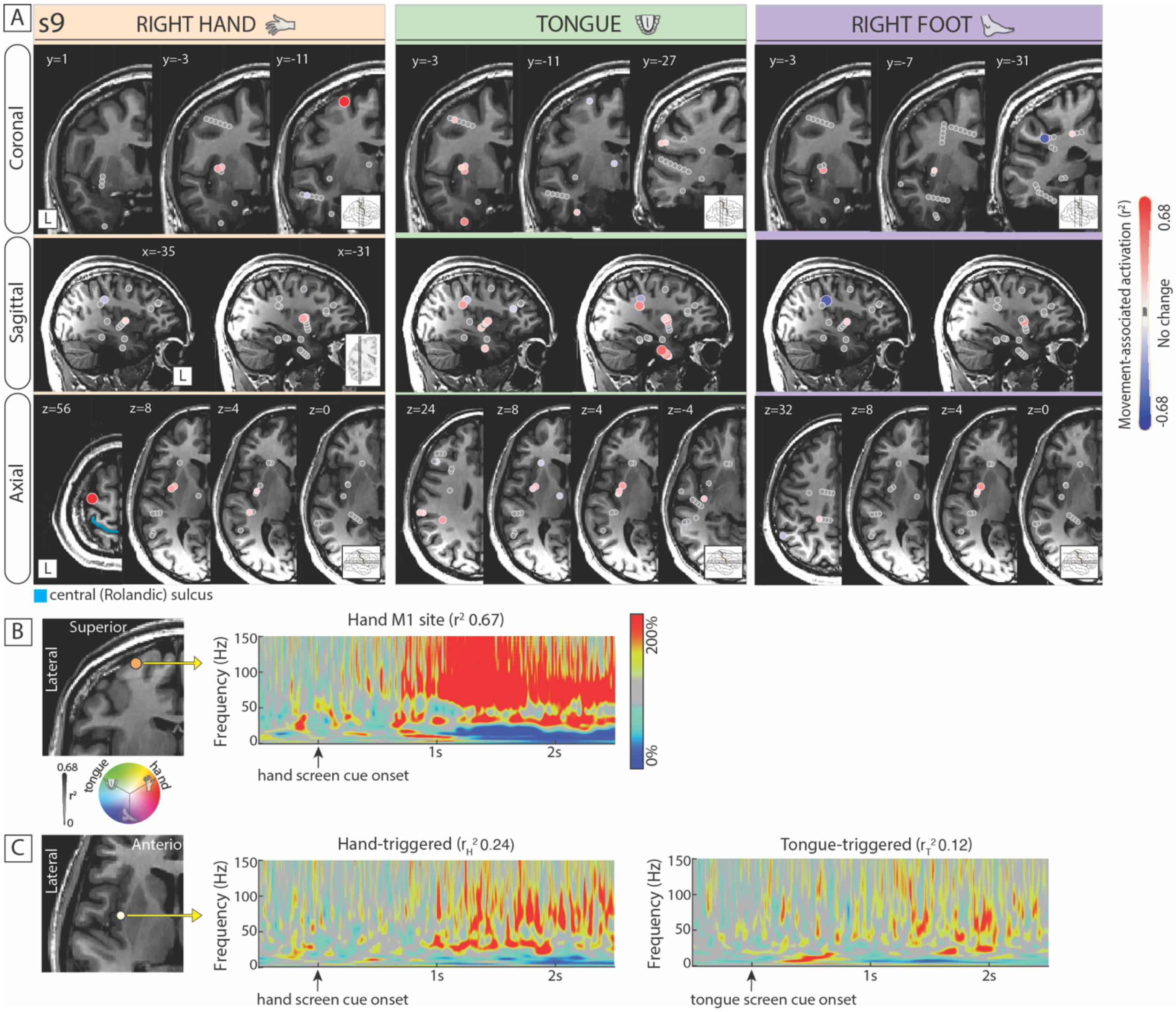


**Suppl. Figure 20.** Posterior insular activation during hand and tongue movement (s9). A) Activation $r^2$ maps for hand (orange), tongue (green) and foot (purple) movement in coronal, sagittal and axial planes (4 mm slices). The peri-insular sulcus is shown in yellow and the central sulcus in blue. (B) Primary motor cortical hand site and its screen cue-triggered average (-0.5 s to 2 s) spectrogram. (C) The posterior insular channel was active during hand and tongue movement.

#### FIGURE S21

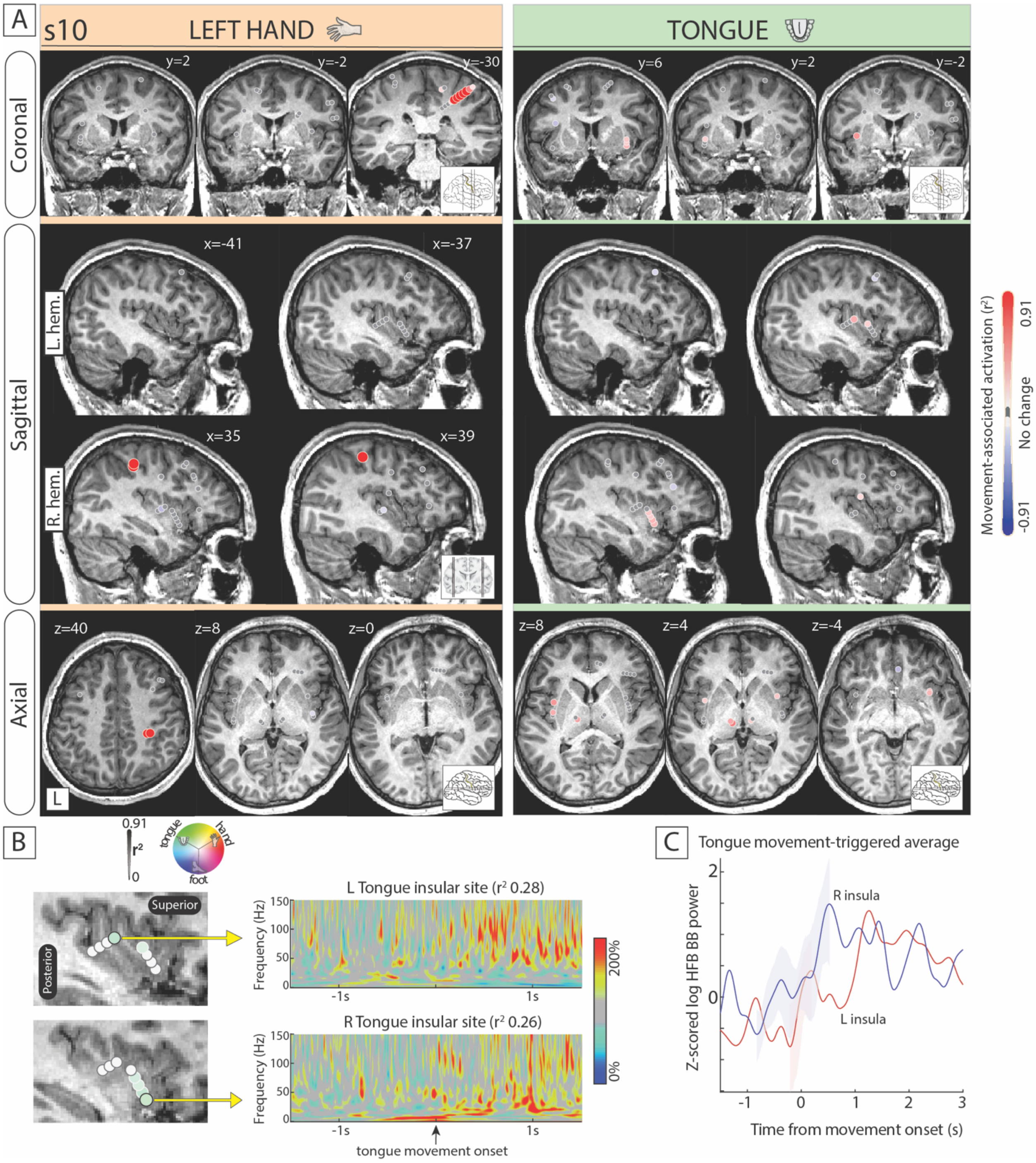


**Suppl. Figure 21.** Bilateral insular activation during tongue movement (s10). (A) Activation $r^2$ maps for hand (orange) and tongue (green) movement in coronal, sagittal and axial planes (4 mm slices). There was no activation during hand or foot movement. (B) Movement-triggered average (-1.5 s to 1.5 s) spectrograms during tongue movement for bilateral tongue-tuned insular sites. (C) Movement-triggered average (-0.5 s to 3 s) high-frequency broadband responses, along with the standard error of the mean during the rising phase, comparing the activity of both tongue-tuned insular sites.

#### FIGURE S22

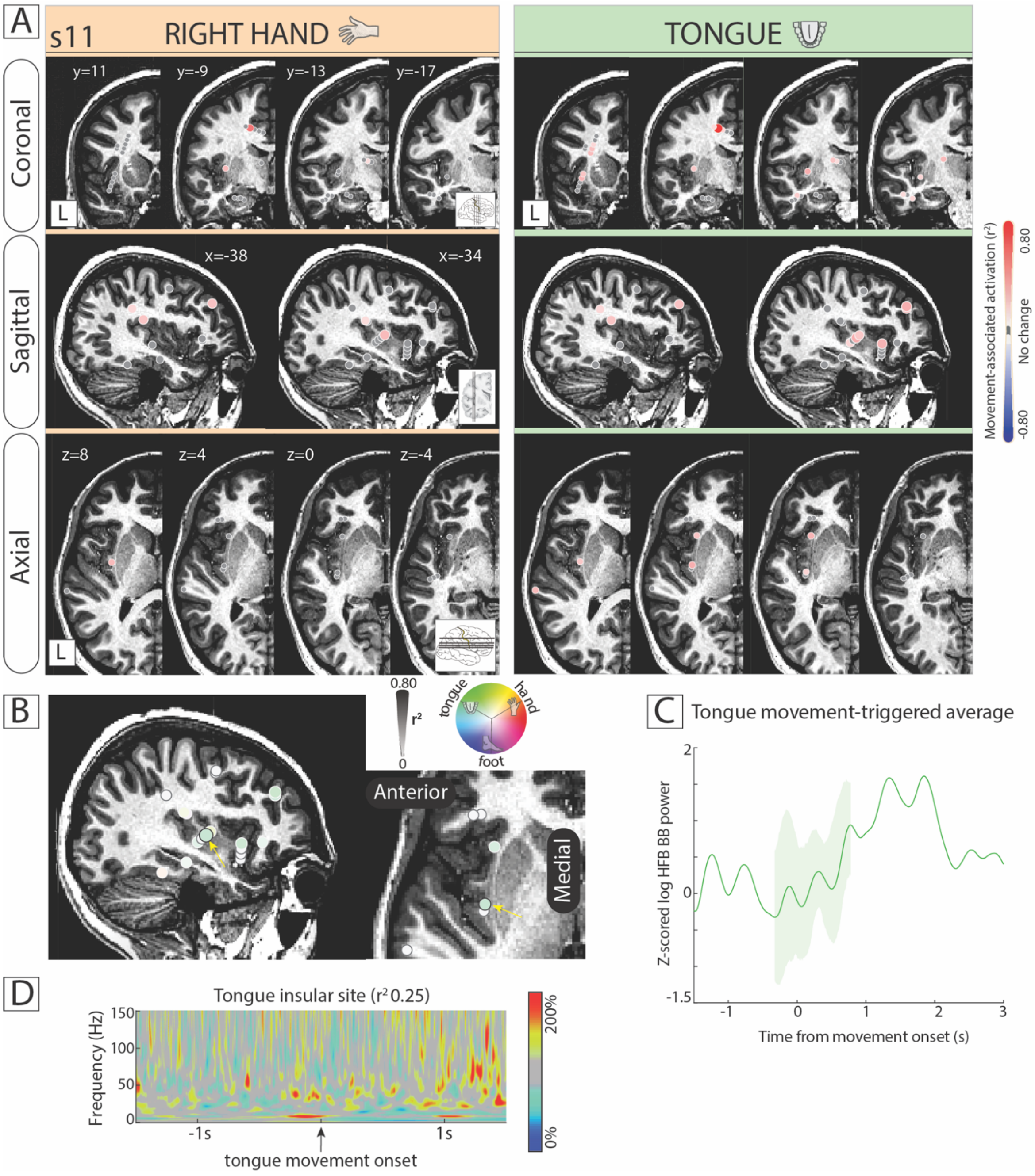


**Suppl. Figure 22.** Somatotopic representation of tongue-movement in the posterior insula (s11). (A) Activation $r^2$ maps for hand (orange) and tongue (green) movement in coronal, sagittal and axial planes (4 mm slices). (B) The tongue-selective insular channels are visualized. (C) Movement-triggered average (-1.5 s to 1.5 s) spectrogram for the tongue-tuned insular site with the highest $r^2$ value. (D) Screen cue-triggered average (-0.5s to 2s) high-frequency broadband response (z-scored) in the same channel. The broadband latency is shown in the inset bar graph.

#### FIGURE S23

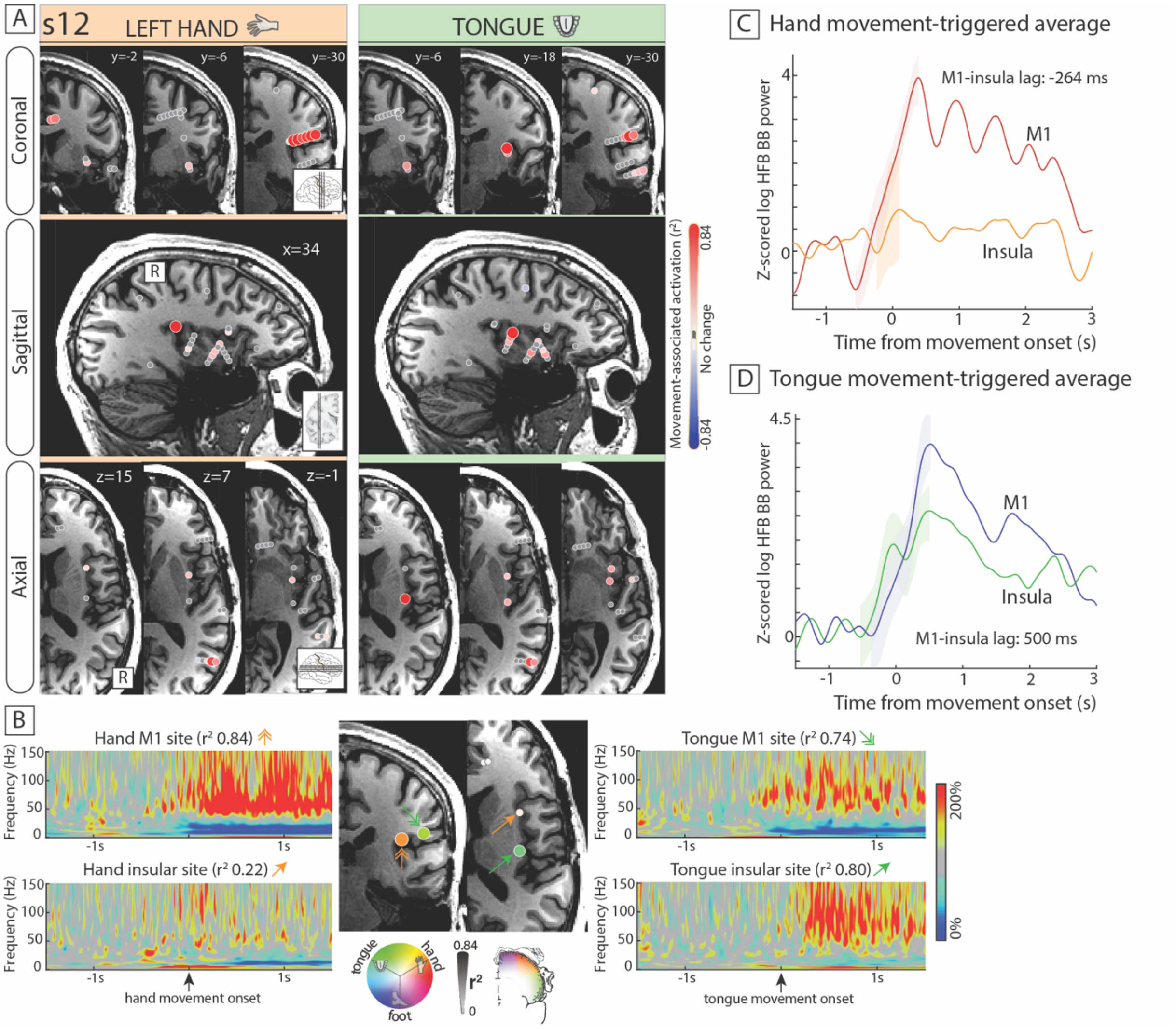


**Suppl. Figure 23.** Somatotopic representation of hand and tongue movement in the insula (s12). (A) Activation $r^2$ maps for hand (orange) and tongue (green) movement in coronal, sagittal and axial planes (4 mm slices). (B) Movement-triggered average (-1.5 s to 1.5 s) spectrograms during hand and tongue movement for the somatotopically-tuned primary motor cortical (double-headed arrows) and insular sites (single-headed arrows). (C, D) Movement-triggered average (-0.5 s to 3 s) high-frequency broadband responses, along with the standard error of the mean during the rising phase, comparing the activity of insular sites to M1 activity and EMG onset.

#### FIGURE S24

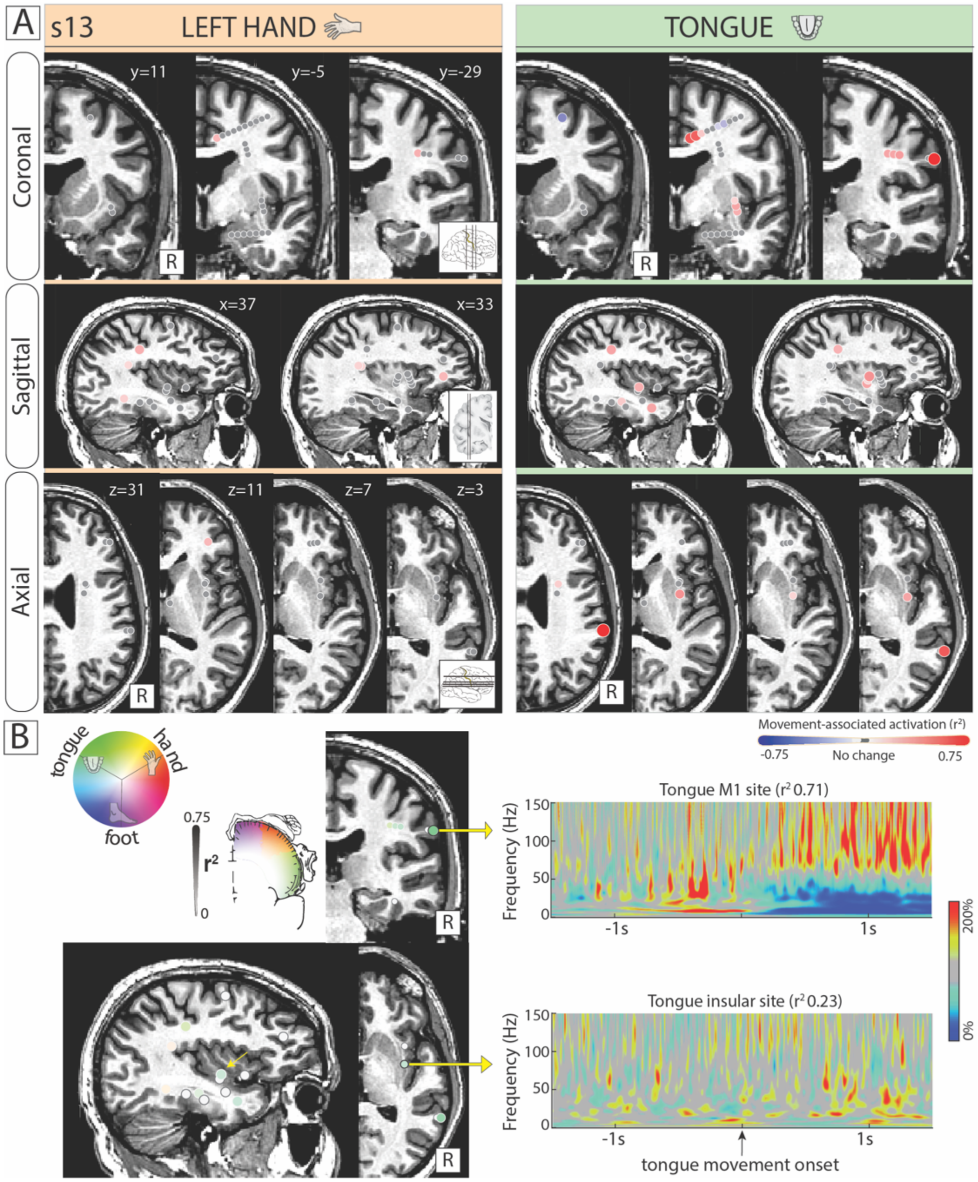


**Suppl. Figure 24.** Somatotopic representation of tongue movement in the posterior insula (s13). Activation $r^2$ maps for hand (orange) and tongue (green) movement in coronal, sagittal and axial planes (4 mm slices). (B) Movement-triggered average (-1.5 s to 1.5 s) spectrograms during tongue movement for the somatotopically-tuned primary motor cortical and insular sites (yellow arrow).

#### FIGURE S25

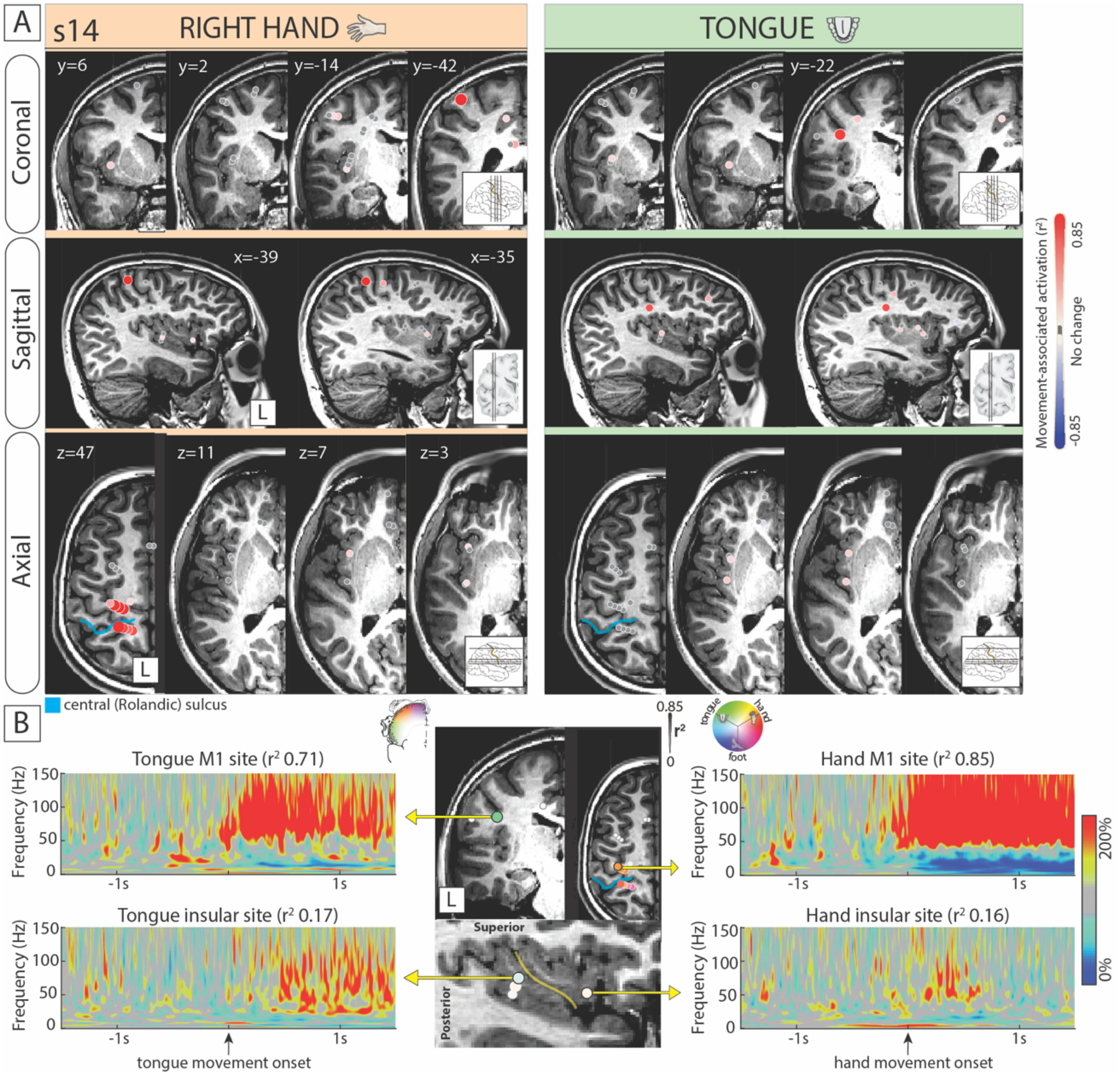


**Suppl. Figure 25.** Insular activation during hand and tongue motor execution (s14). (A) Activation $r^2$ maps for hand (orange) and tongue (green) movement in coronal, sagittal and axial planes (4 mm slices). The central sulcus in blue. (B) Movement-triggered average (-1.5 s to 1.5 s) spectrograms during hand and tongue movement for the somatotopically-tuned primary motor cortical and insular sites. The central sulcus is shown in blue.

#### FIGURE S26

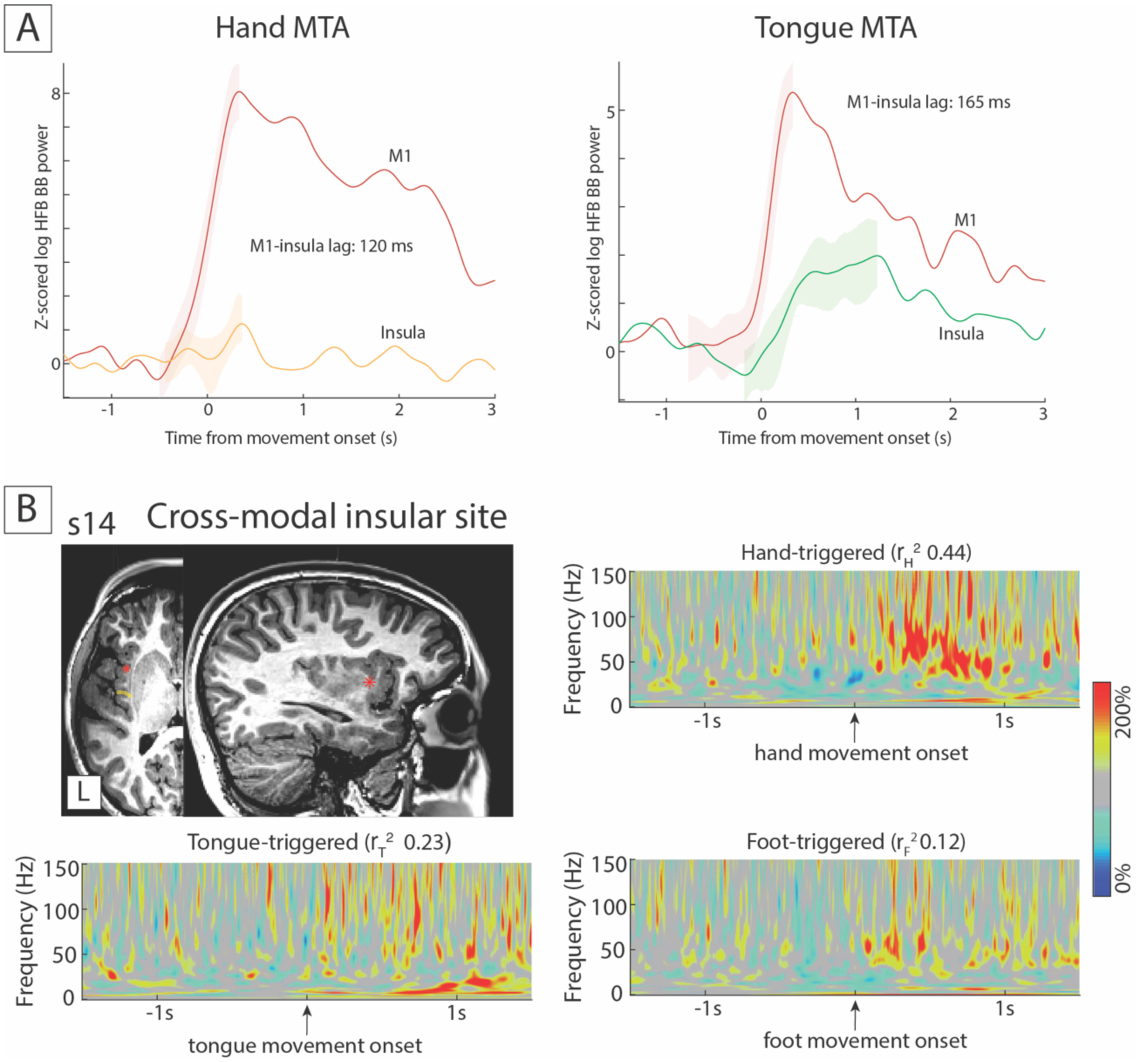


**Suppl. Figure 26.** (A) Hand and tongue movement triggered average (-0.5 s to 3 s) high-frequency broadband responses, along with the standard error of the mean during the rising phase, comparing the activity of insular sites to M1 activity. (B) Movement-triggered average time-frequency plots (-1.5 s to 1.5 s) during hand, tongue and foot movement in the cross-modal channel in the anterior-dorsal insula.

#### FIGURE S27

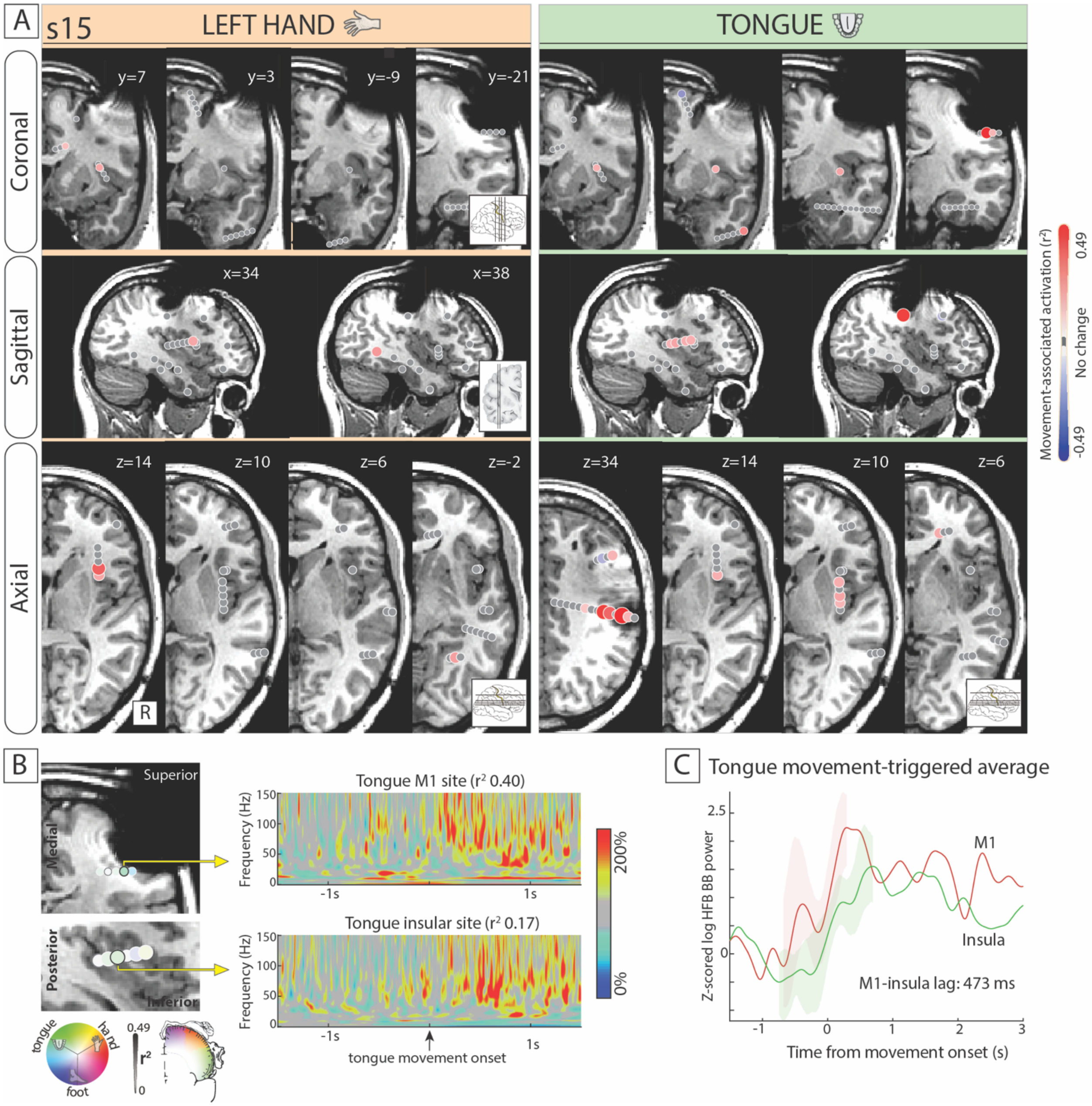


**Suppl. Figure 27.** Tongue insular motor representation in subject 15. Please note right frontal artifact due to existing ventriculo-peritoneal shunt from young age for hydrocephalus. (A) Activation $r^2$ maps for hand (orange) and tongue (green) movement in coronal, sagittal and axial planes (4 mm slices). (B) Tongue-selective channels in the primary motor cortex and posterior insula and their respective movement-triggered average (-0.5 s to 1.5 s) spectrograms. (C) Movement-triggered average (-0.5 s to 3 s) high-frequency broadband responses, along with the standard error of the mean during the rising phase, comparing the activity of the tongue insular site to M1 activity.

### FIGURE S28

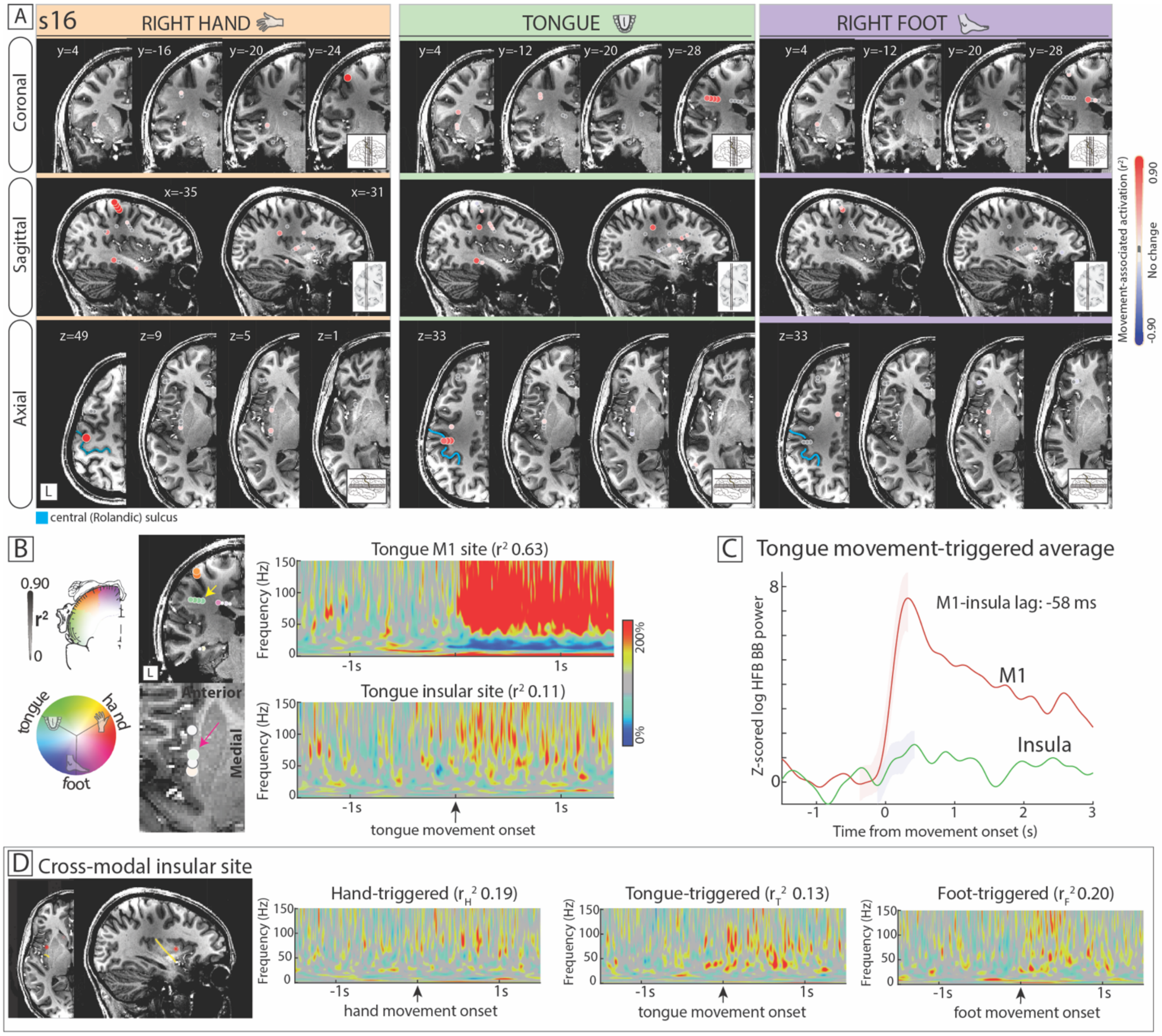


**Suppl. Figure 28.** Insular motor representation in a case with tongue-tuned and inter-effector insular channels (s16). (A) Activation $r^2$ maps for hand (orange), tongue (green) and foot (purple) movement in coronal, sagittal and axial planes (4 mm slices). The central sulcus in blue. (B) Movement-triggered average (-0.5 s to 1.5 s) spectrograms for the tongue-tuned sites in the primary motor cortex (yellow arrow) and posterior insula (green arrow). (C) Movement-triggered average (-0.5 s to 3 s) high-frequency broadband responses, along with the standard error of the mean during the rising phase, comparing the activity of the tongue insular sites to M1 activity. (D) Cross-modal insular channel in the anterior-dorsal insula (central sulcus in yellow) and the movement-triggered average spectrogram for each movement type.

#### FIGURE S29

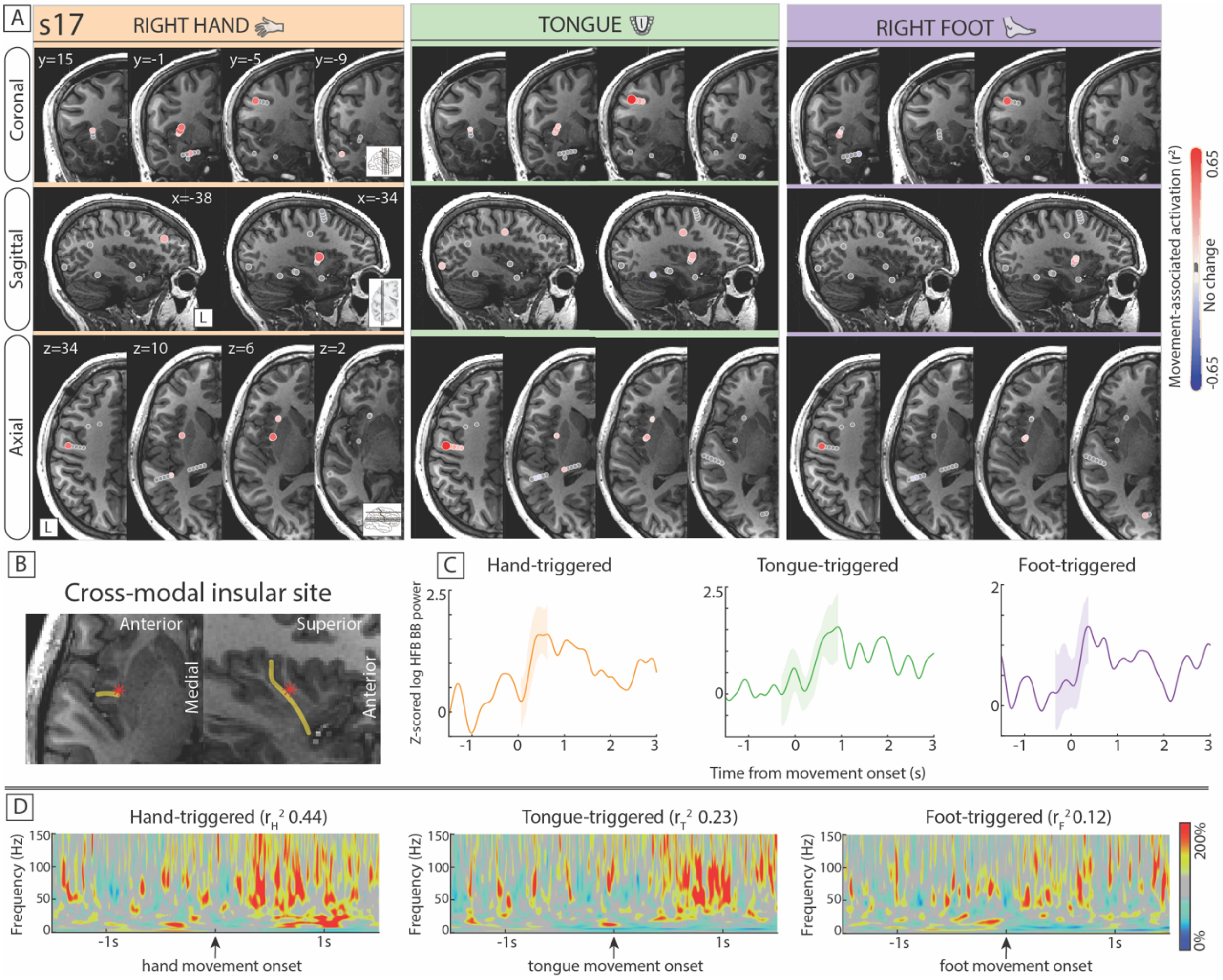


**Suppl Figure 29.** Shared, inter-effector insular motor representation in subject 17. (A) Activation $r^2$ maps (s3) for hand (orange), tongue (green) and foot (purple) movement in coronal, sagittal and axial planes (4 mm slices). (B) The crossmodal motor site was identified just anterior to the central sulcus of the insula. We did not identify hand- or tongue-selective channels in this patient. (C) Movement-triggered average (-0.5 s to 2 s) high-frequency Movement-triggered average (-0.5 s to 3 s) high-frequency broadband responses, along with the standard error of the mean during the rising phase, for each movement type. (D) Movement-triggered average (-0.5 s to 1.5 s) spectrograms for each movement type in the same channel.
